# Rapid evaporative ionisation mass spectrometry (REIMS): A potential and rapid tool for the identification of insecticide resistance in mosquito larvae

**DOI:** 10.1101/2022.02.10.479854

**Authors:** Jasmine Morgan, J. Enrique Salcedo-Sora, Iris Wagner, Robert J Beynon, Omar Triana-Chavez, Clare Strode

## Abstract

Insecticide resistance is a significant challenge facing the successful control of mosquito vectors globally. Bioassays are currently the only method for phenotyping resistance. They require large numbers of mosquitoes for testing, the availability of a susceptible comparator strain and often insectary facilities. This study aimed to trial the novel use of rapid evaporative ionisation mass spectrometry (REIMS) for the identification of insecticide resistance in mosquitoes. No sample preparation is required for REIMS and analysis can be rapidly conducted within hours. Temephos resistant *Aedes aegypti* (Linnaeus) larvae from Cúcuta, Colombia and temephos susceptible larvae from two origins (Bello, Colombia, and the lab reference strain New Orleans) were analysed using REIMS. We tested the ability of REIMS to differentiate three relevant variants: population source, lab versus field origin and response to insecticide. The classification of these data was undertaken using linear discriminant analysis (LDA) and random forest. Classification models built using REIMS data were able to differentiate between *Ae. aegypti* larvae from different populations with 82% (± 0.01) accuracy, between mosquitoes of field and lab origin with 89% (± 0.01) accuracy and between susceptible and resistant larvae with 85% (± 0.01) accuracy. LDA classifiers had higher efficiency than random forest with this data set. The high accuracy observed here identifies REIMS as a potential new tool for rapid identification of resistance in mosquitoes. We argue that REIMS and similar modern phenotyping alternatives should complement existing insecticide resistance management tools.

## Introduction

Insecticide resistance is one of the most significant challenges posed to mosquito control programmes. The control of mosquito vectors, including *Aedes aegypti* (Linnaeus) the principal vector for the dengue, Zika and chikungunya viruses, relies heavily on the use of insecticides to reduce disease burden. There are only four insecticides classes which are licensed for use in public health: organophosphates, organochlorines, pyrethroids and carbamates. Resistance has now been reported in *Ae. aegypti* to all four of these chemical classes (Ranson et al. 2010, Vontas et al. 2012, Moyes et al. 2017). Insecticide resistance in *Ae. aegypti* is also spread worldwide with reports in South America (Guedes et al. 2020), North America (Marcombe et al. 2014), Asia (Amelia-Yap et al. 2018), Europe (Seixas et al. 2017), Africa (Weetman et al. 2018), and Oceania (Demok et al. 2019). This trend is compromising effective vector control (Viana-Medeiros et al. 2007, Bisset et al. 2011, Marcombe et al. 2011).

Insecticide resistance management (IRM) which aims to prevent, slow, or reverse the emergence of resistance is therefore crucial for sustainable vector control. The first step in IRM is to monitor local populations for the development of insecticide resistance whilst establishing its impact on effective vector control (Dusfour et al. 2019). Current methods for resistance monitoring include bioassays, biochemical assays, and molecular testing. Biochemical assays and molecular testing are used to identify the specific mechanisms responsible for insecticide resistance, allowing for appropriate IRM strategies to be implemented (Hemingway et al. 2013). However, insecticide bioassays (e.g. WHO tube and CDC bottle assays) are the only current method for identifying (phenotyping) resistance in mosquitoes. Bioassays have low sensitivity, lengthy completion times (24 hours) and often only detect high levels of resistance which maybe too late for alternative measures to be deployed (Dusfour et al. 2019). Other limitations include the requirement of large numbers of individual mosquitoes, and the availability of a comparable susceptible strain (World Health Organization (WHO) 2016). Alternative phenotyping methods that can surpass those limitations are necessary.

Rapid evaporative ionisation mass spectrometry (REIMS) is a relatively new technology which provides a rapid method of mass spectrometry without the need for any sample preparation. Samples are burned by diathermy and the resultant aerosols are collected, ionized, and analysed by mass spectrometry (Schäfer et al. 2009, Balog et al. 2010, 2013, 2015). The spectra, collected in negative ion mode, largely reflect the lipid composition of the sample, and is collected over a wide range of m/z values. The spectra are then are discretised by binning, creating a data matrix that is further processed by dimension reduction and classification (Balog et al. 2010). The potential applications of REIMS are vast with its previous successful applications including distinguishing cancerous tissue from healthy tissue (Alexander et al. 2017, St John et al. 2017, Phelps et al. 2018), authentication of food products (Balog et al. 2016, Black et al. 2017, Verplanken et al. 2017, Guitton et al. 2018, Rigano et al. 2019), microbial species identification (Strittmatter et al. 2013, 2014), monitoring of bacterial growth and recombinant protein expression (Sarsby et al. 2021), and the identification of rodent species and sex from faecal matter (Davidson et al. 2019). REIMS has also been shown to be a highly effective method for species and sex determination in *Drosophila* adults and larvae (Wagner et al. 2020).

Here we present a proof-of-concept for the novel use of REIMS as a rapid tool for the identification of insecticide resistance in *Ae. aegypti* larvae. We analysed three *Ae. aegypti* populations, previously profiled for susceptibility to the larvicide temephos (Morgan et al. 2021): a resistant population originating from field collected mosquitoes from Cúcuta (Colombia) and two susceptible populations, one field originating population from Bello (Colombia) and a susceptible laboratory reference strain, New Orleans. The results demonstrate the potential of REIMS for phenotyping insecticide resistant mosquitoes with relevant discriminatory power and faster and less labour-intensive methods which may be used to complement existing IRM strategies.

## Materials and methods

### Mosquito samples and rearing

*Aedes aegypti* larvae from three populations previously tested for susceptibility to temephos (Morgan et al. 2021) were used in this study. Two field populations were used, one temephos resistant (field resistant (FR) and one susceptible (field susceptible (FS)), the susceptible *Ae. aegypti* laboratory strain New Orleans (lab susceptible (LS)) was also used (Fig.1). *Ae. aegypti* were reared to fourth instar larvae following a standard rearing protocol and under standard conditions within Edge Hill University Vector Research Group insectaries. Standard conditions were 27°C and 70% relative humidity with an 11-hour day/night cycle with 60-minute dawn/dusk simulation periods, using a lighting system of 4× Osram Dulux 26W 840 lights. Eggs were submerged in a hatching broth of 350 ml dH_2_O, 0.125 g nutrient broth (Sigma-Aldrich, Dorset, UK) and 0.025 g brewer’s yeast (Holland & Barrett, Ormskirk, UK) for 48 hours (Zheng et al. 2015). Once hatched, larvae were reared at a density of 0.5 larva/ml in dH_2_O and fed ground fish food (AQUARIAN® advanced nutrition) at increasing quantities per day (day 3 = 0.08 mg/larva, day 4 = 0.16 mg/larva, day 5 = 0.31 mg/larva, day 6 = 0 mg/larva) (Carvalho et al. 2014). For each experimental group (FR, FS, LS) four biological replicates were conducted, using eggs from different females each submerged on different days. Seven days after egg submission larvae were removed and stored at -20°C until REIMS analysis. The storage period ranged from 32-36 weeks (Table 1). The number of larvae analysed per biological replicate ranged from 8-15 with a total of 42-51 larvae per experimental group (Table 1).

**Fig. 1:**
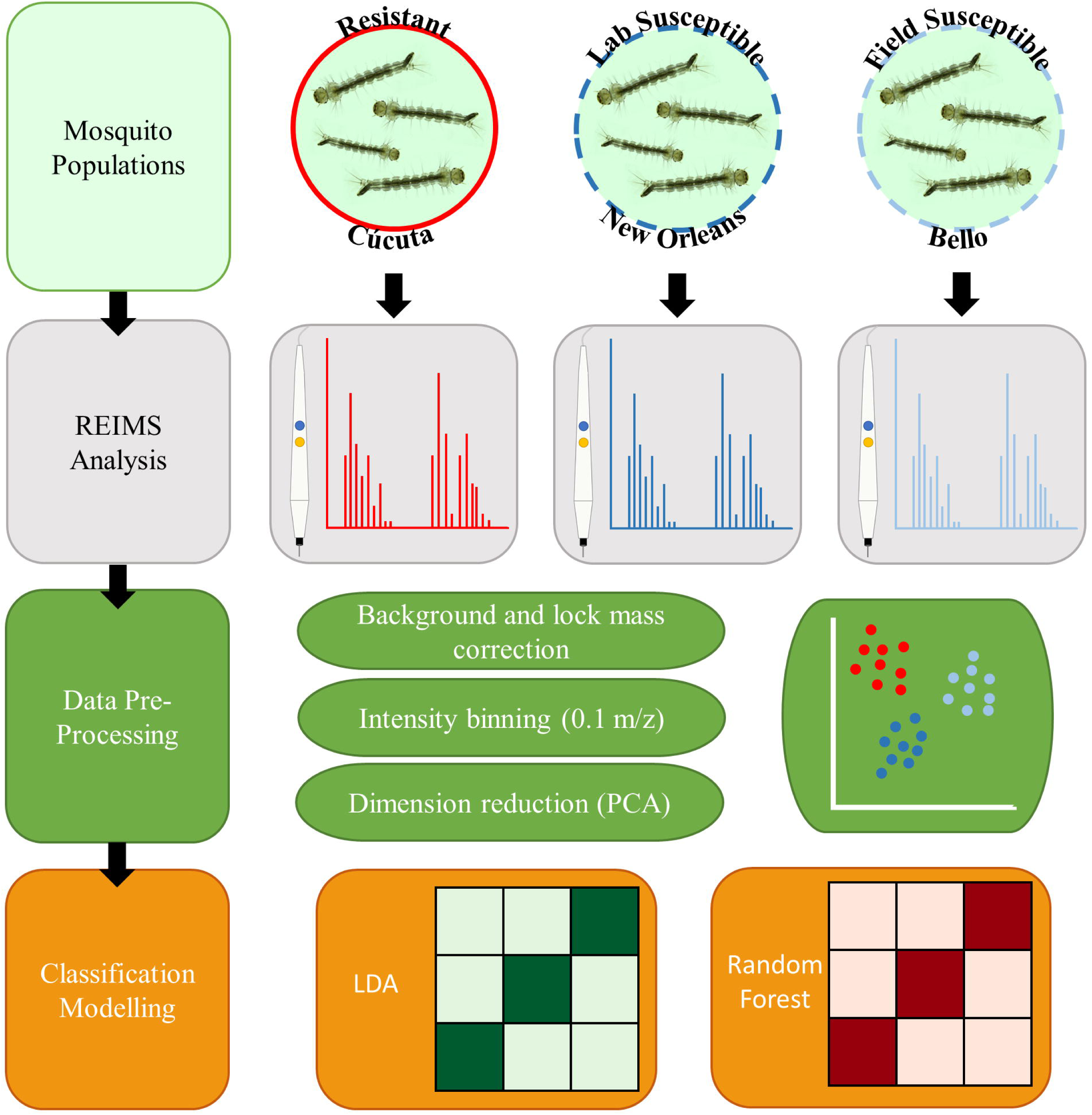
Block diagram of the experimental approach. This study utilised insecticide resistant and susceptible larvae of the mosquito *Ae. aegypti*. The resistant larvae originated from Cúcuta, Colombia and the susceptible larvae had dual origin, field samples from Bello, Colombia (Field Susceptible) and the New Orleans lab strain (Lab Susceptible). Individual larvae from each experimental group were analysed using REIMS to acquire individual mass spectra for each sample. The data acquired through REIMS was background and lock mass corrected and binned into 0.1 m/z groups. Dimension reduction was conducted using PCA before LDA and random forest classification model building and testing.

**Table 1:** Summary data of the *Ae. aegypti* samples analysed via REIMS. Time larvae stored at -20°C in weeks for each replicate and the number of larvae analysed in each replicate and the total number for each experimental group (*n*).

| Population | Replicate | Storage Weeks | *n* |
| --- | --- | --- | --- |
| Lab<br>Susceptible | 1 | 36 | 8 |
|  | 2 | 36 | 15 |
|  | 3 | 36 | 13 |
|  | 4 | 32 | 15 |
|  | Total | 32-36 | 51 |
| Field<br>Susceptible | 1 | 32 | 12 |
|  | 2 | 34 | 13 |
|  | 3 | 33 | 13 |
|  | 4 | 32 | 13 |
|  | Total | 32-34 | 51 |
| Field<br>Resistant | 1 | 36 | 9 |
|  | 2 | 32 | 14 |
|  | 3 | 36 | 10 |
|  | 4 | 36 | 9 |
|  | Total | 32-36 | 42 |

### Rapid evaporative ionisation mass spectrometry analysis

Rapid evaporative ionisation mass spectrometry analysis was conducted following the detailed methods outlined by Wagner *et al*. (2020). Larvae were burned using a monopolar electrosurgical pencil (Erbe Medical UK Ltd, Leeds); the electric current was provided to the pencil by a VIO 50 C electrosurgical generator, a black conductive rubber mat acted as the counter electrode to enable the flow of electricity through the sample. The entire biomass of each larva was burned, and the aerosols produced were aspirated through tubing attached to the pencil into the REIMS source using a nitrogen powered venturi valve. Leucine enkephalin (Waters, UK) in propan-2-ol (CHROMASOLV, Honeywell Riedel-de-Haën) was used as a lock mass solution and continuously introduced via a whistle in the venturi tube at a flow rate of 30 µl min−1. REIMS was conducted using a Synapt G2Si instrument ion mobility equipped quadrupole time of flight mass spectrometer (Waters, UK). A heated impactor (Kanthal metal coil at 900°C) within the REIMS source was used to decluster the ionized particles. Mass spectra were acquired in negative ion mode at a rate of 1 scan per second over a mass/charge range of *m/z* 50–1200. All larvae were analysed in a single day in a random order created by a random number generator within Microsoft excel.

### Data analysis

The raw data files were imported into the Offline Model Builder software (OMB-1.1.28; Waters Research Centre, Hungary). Each data file/sample contains the burn event of only one larva, therefore the option to create one spectrum per sample was selected. The background was subtracted, and the spectra corrected using the lock mass (leucine enkephalin, m/z 554.26). The normalised intensities were then binned into 0.1 m/z wide groups. The binned mass spectra data were then imported into R (version 3.6.3) (R Core Team 2020) for further analysis.

Dimension reduction was carried out by principal components analysis (PCA) using the R package factoextra (version 1.0.7) (Kassambara and Mundt 2020). Different numbers of principal components were then extracted (10,20,40,60,80,100) and used for classification of samples into categories: population, population type and resistance status. Classification was conducted using two different model types; linear discriminant analysis (LDA) and random forest (RF), with the data randomly split into 70% training data and 30% test data. Each model was built using variable numbers of principal components (PCs) extracted using PCA and the most accurate model selected and used for analysis. LDA models with varying numbers of PCs were built using the R package MASS (version 7.3.53) (W. N. Venables and B. D. Ripley 2002), model validation was conducted by plotting receiver operating characteristic curves (ROC) and selecting the model with the highest area under ROC curve (AUC) (Supp Fig.S2-11). Random forest models were validated using the R package caret (version 6.0.88) (Kuhn 2021) to select the model with optimum PCs, number of variables available for splitting at each tree node (mtry) and tree number. The random forest models with the highest overall accuracy following building in caret were selected for use in the analysis with models built using the R package randomForest (version 4.6.14) (Liaw and Wiener 2002). Random under sampling in the caret package was used to balance classes prior to RF analysis as this showed increase in overall model performance. Class imbalance did not affect performance of LDA models, as no difference in classification accuracy was observed between the different groups within the models, therefore no over or under sampling was required. LDA and RF models with parameters as selected by model validation were each ran 20 times using a different random split of test (30%) and training (70%) data. The model statistics: percentage accuracy, standard error of means (SEM) and range, were then averaged across all 20 replicates. LDA following PCA was also used to visualise the separation of samples, plots were created using the R packages ggplot2 (version 3.3.2) (Wickham 2016) and ggpubr (version 0.4.0) (Kassambara 2020).

The experimental design is outlined in Fig.1. A code for analysing REIMS data using LDA and random forest classification models which can be applied to other similar datasets is available in Supplementary File 1. All raw data files are available in the MetaboLights database under the accession number MTBLS4129. The data matrix, created in OMB and used for subsequent analysis in R is available in Supplementary table 1.

## Results

### Population source

Visualisation of the data, following PCA-LDA analysis showed a clear discrimination between *Ae. aegypti* larvae from different geographical origins (Fig.2A). All three populations; field susceptible, field resistant and lab susceptible were separated in linear discriminant one whilst the field resistant population separated from the two susceptible populations in linear discriminant 2, thus demonstrating that LD1 is representative of population and LD2 of resistance to insecticide. A PCA-LDA conducted on the data with randomly assigned classifications showed no separation (Supp Fig.S1) demonstrating that the observed separation of classifications is due to variations between populations and not due to chance. The LDA model built using the REIMS data was able to correctly classify 82% (± 0.01) of *Ae. aegypti* larvae into the correct population (Fig.2B). The lab susceptible population had the highest accuracy (90% ± 2.0) and had the largest sample number whilst the population with the lowest sample number, field resistant, had the lowest accuracy (77% ± 2.2). When classification was conducted using a random forest model accuracy was lower, but the model was still able to correctly assign 76% of individual *Ae. aegypti* larvae to the correct population (Fig.2C).

**Fig. 2:**
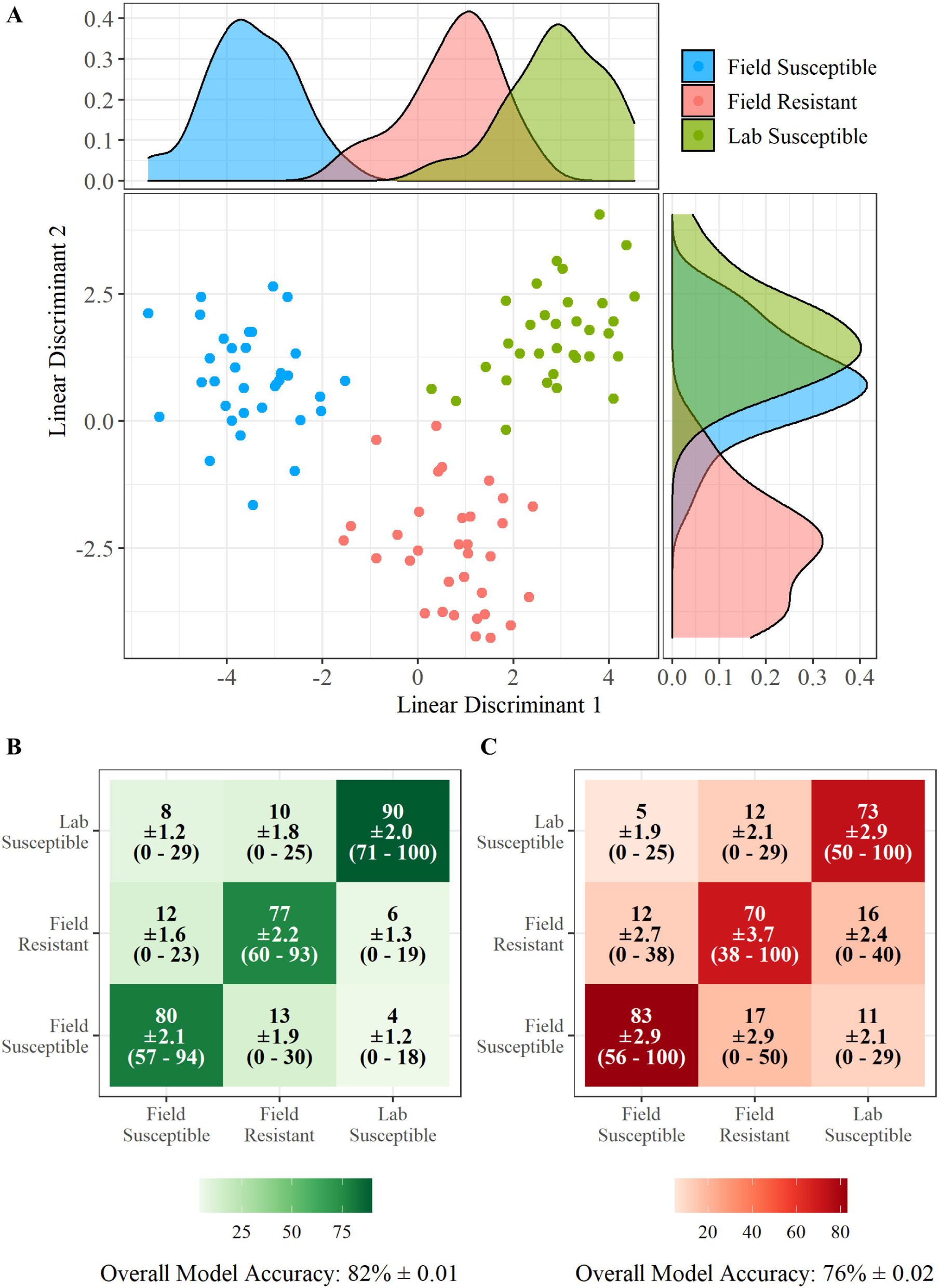
REIMS discrimination of *Ae. aegypti* samples by population. Combined PCA-LDA separation of the three *Ae. aegypti* populations using REIMS mass spectra (A). Dimension reduction was conducted using principal components analysis (PCA), 40 principal components were selected for linear discriminant analysis (LDA). The number of PCs was determined by selecting the model with the lowest area under the ROC curve (AUC) (Supp Fig.S2). Separation is shown in both linear discriminant one and linear discriminant 2. All populations separated in linear discriminant 1 whilst field resistant separated from the two susceptible populations in LD2. Classification of samples into population using PCA-LDA (B) and random forest models (C), showing percentage of samples classified to each group, standard error of the mean (SEM) and the percentage range across all replicates. Models were built and tested 20 times each with a different set of training (70%) and test (30%) data. Accuracy percentages, SEM and range were averaged across all 20 replicates. The PCA-LDA classification model had a higher accuracy (82% ± 0.01) than the random forest model (76% ± 0.02), correctly assigning 82% of individuals to their respective population. Random forest models were built using 20 PCs to obtain the highest accuracy of models tested (Supp Fig.S3).

### Population type (lab and field)

A clear separation is observed when *Ae. aegypti* larvae from field origin are compared to larvae from a standard laboratory reference strain using PCA-LDA (Fig.3A & B). The classification models had high accuracy with 89% (± 0.01) of individual larvae classified to the correct population type with the PCA-LDA model (Fig.3C) and 83% (± 0.01) correctly classified by random forest (Fig.3D). Larvae from field origin had higher classification accuracy (86% ± 1.8) than those of lab origin (80% ± 2.4) when the RF model was used. When the LDA model was used the accuracy was similar for both groups (Field = 90% ± 0.8, Lab = 89 ± 2.0).

**Fig. 3:**
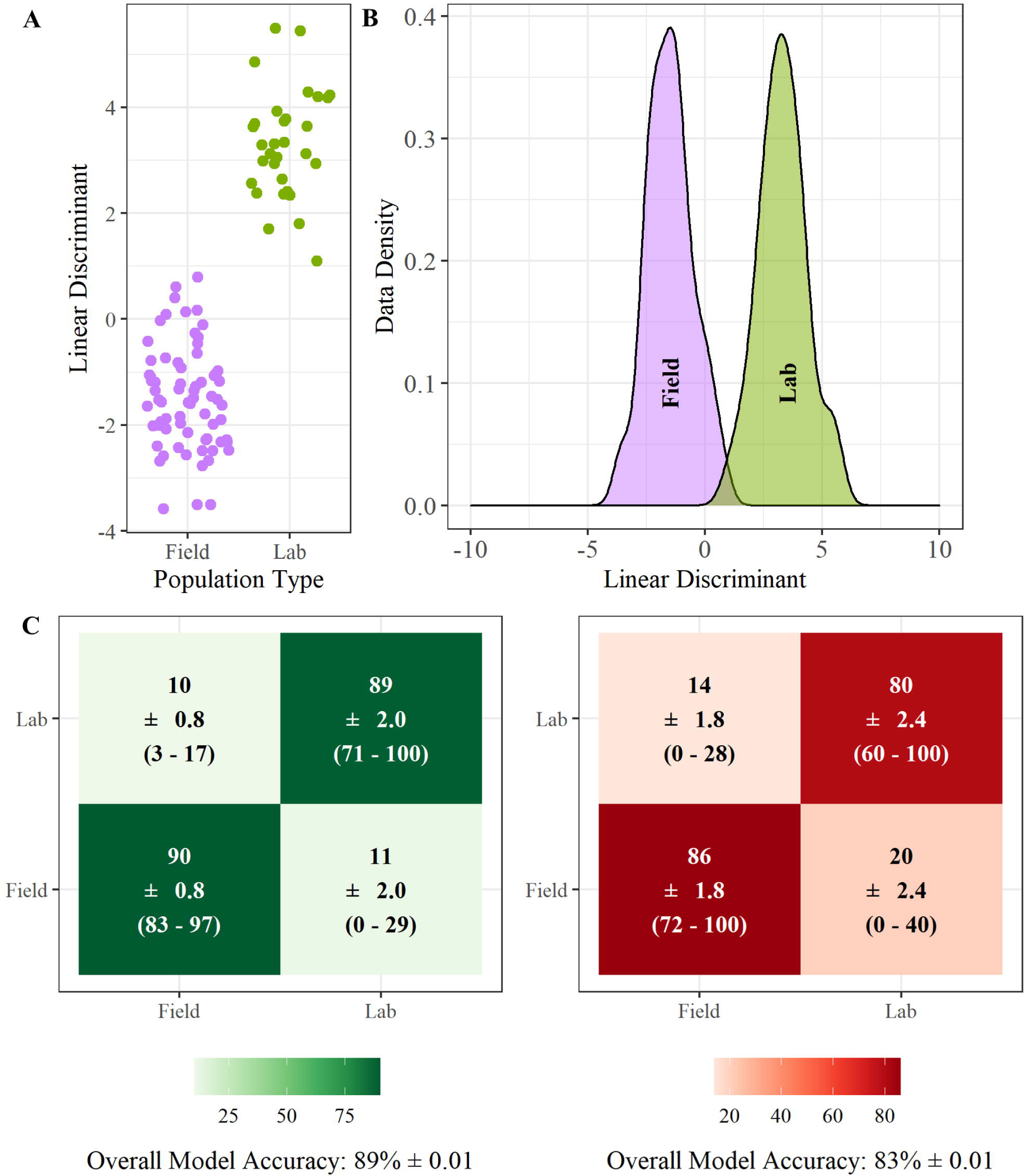
REIMS discrimination of *Ae. aegypti* by population type (lab and field). Combined PCA-LDA separation of lab and field *Ae. aegypti* populations using REIMS mass spectra (A & B). Dimension reduction was conducted using principal components analysis (PCA), 40 principal components were selected for linear discriminant analysis (LDA). The number of PCs was determined by selecting the model with the lowest area under the ROC curve (AUC) (Supp Fig.S4). Classification of samples into resistance status using PCA-LDA (B) and random forest models (D), showing percentage of samples classified to each group, standard error of the mean (SEM) and the percentage range across all replicates. Models were built and tested 20 times each with a different set of training (70%) and test (30%) data. Accuracy percentages, SEM and range were averaged across all 20 replicates. The LDA-PCA classification model had a higher accuracy (89% ± 0.01) than the random forest model (83% ± 0.02), correctly assigning 89% of individuals to their respective resistance status. Random forest models were built using 20 PCs to obtain the highest accuracy of models tested (Supp Fig.S5).

### Insecticide sensitivity profile

Analysis of the REIMS data was also conducted to investigate the potential for determination between insecticide resistant and susceptible *Ae. aegypti* larvae (Fig.4). PCA-LDA classification models show 85% (± 0.01) accuracy in assigning larvae to the correct resistance status, with 75% (± 2.8) of temephos resistant larvae being correctly assigned (Fig.4c). The classification accuracy was higher for susceptible individuals (89% ± 1.1), this is likely due to the larger sample size of susceptible individuals available for training the model (Fig.4C). Whilst the random forest classification model was less accurate it still had a correct classification rate of 78% (± 0.02) correctly classifying 73% (± 3.3) of resistant individuals and 79% of susceptible individuals (Fig.4D).

**Fig. 4:**
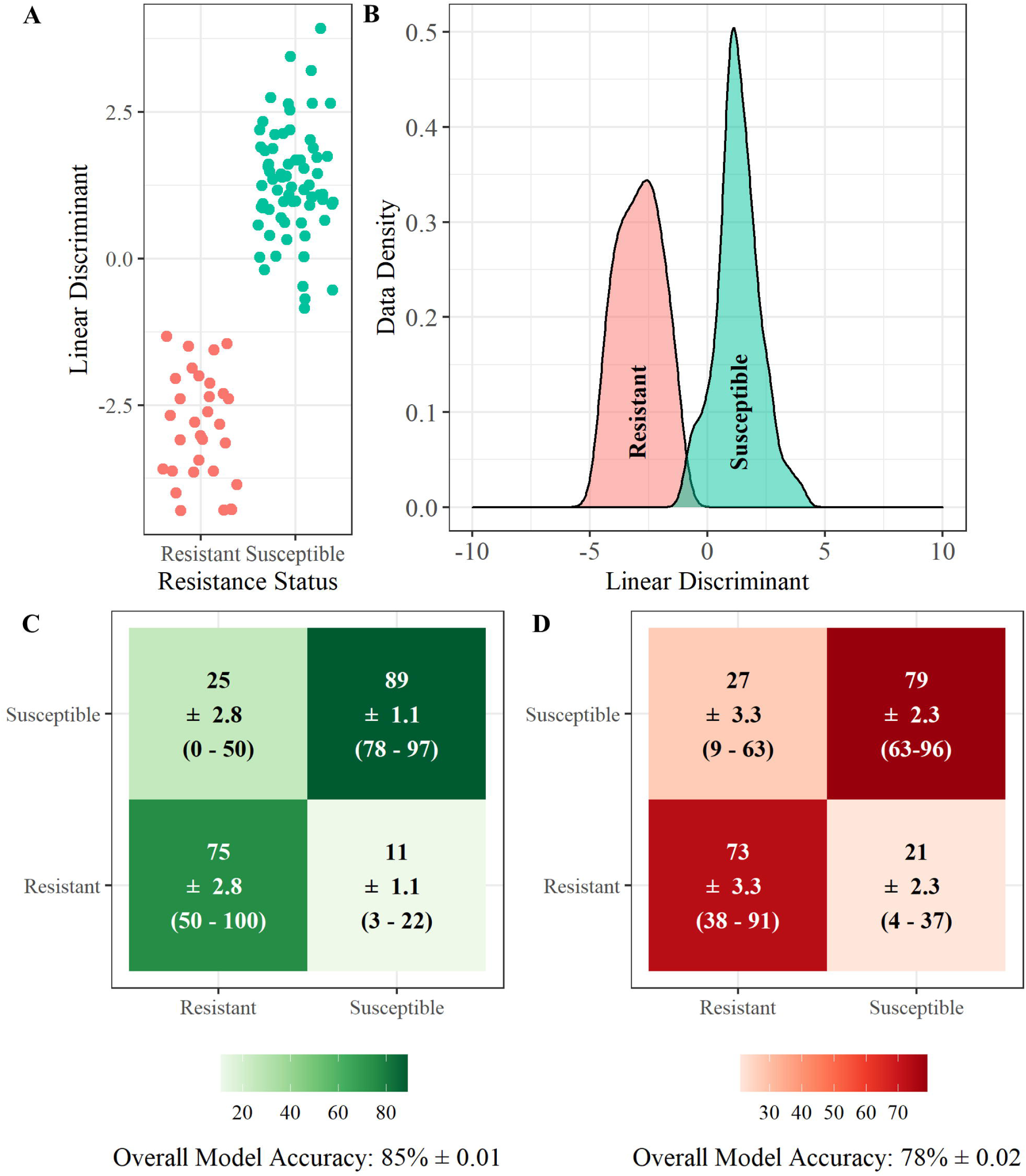
REIMS discrimination of resistant and susceptible *Ae. aegypti*. Combined PCA-LDA separation of resistant and susceptible *Ae. aegypti* populations using REIMS mass spectra (A & B). Dimension reduction was conducted using principal components analysis (PCA), 40 principal components were selected for linear discriminant analysis (LDA). The number of PCs was determined by selecting the model with the lowest area under the ROC curve (AUC) (Supp Fig.S6). Classification of samples into resistance status using PCA-LDA (C) and random forest models (D), showing percentage of samples classified to each group, standard error of the mean (SEM) and the percentage range across all replicates. Models were built and tested 20 times each with a different set of training (70%) and test (30%) data. Accuracy percentages, SEM and range were averaged across all 20 replicates. The LDA-PCA classification model had a higher accuracy (85% ± 0.01) than the random forest model (78% ± 0.02), correctly assigning 85% of individuals to their respective resistance status. Random forest models were built using 20 PCs to obtain the highest accuracy of models tested (Supp Fig.S7).

A similar classification accuracy is achieved when field resistant larvae are compared only to susceptible larvae from a laboratory strain (Fig.5) as when field resistance larvae are compared to susceptible larvae from field origin (Fig.6). When only a field susceptible comparator strain is used the classification accuracy was 88% (± 0.01) using LDA (Fig.6C) and 84% (± 0.02) using RF (Fig.6D). When only a lab susceptible comparator strain is used the classification accuracy was similar with accuracies of 87% with LDA (Fig.5C) and 82% with RF (Fig.5D). The similarity in classification accuracy observed here demonstrates that a field equivalent susceptible strain may not be necessary for identification of insecticide resistance in field *Ae. aegypti* larvae using this method, which is beneficial with the decreasing availability of field relevant susceptible populations.

**Fig. 5:**
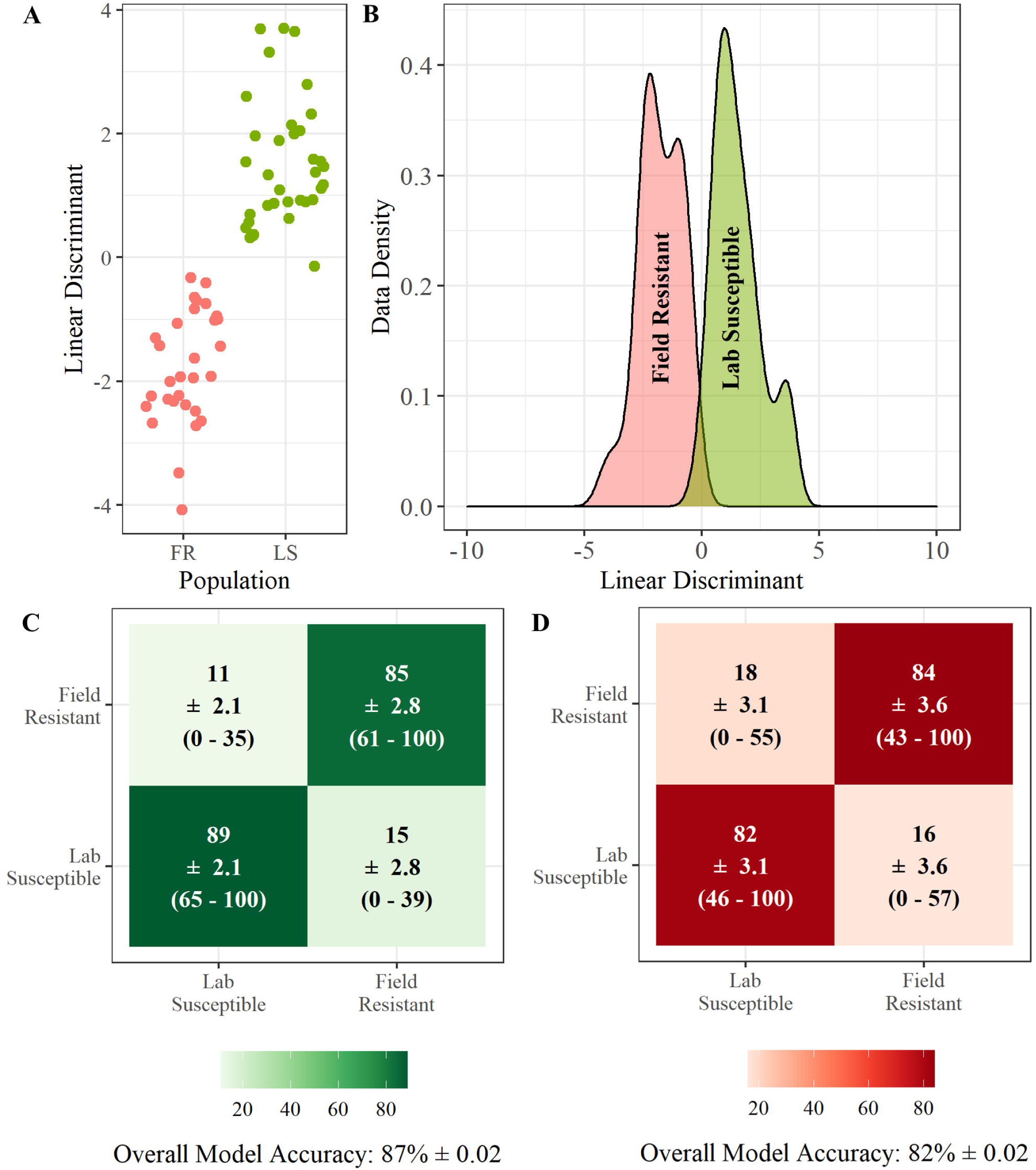
REIMS discrimination of field resistant and lab susceptible *Ae. aegypti* larvae. Combined PCA-LDA separation of the resistant and lab susceptible populations using REIMS mass spectra (A & B). Dimension reduction was conducted using principal components analysis (PCA), 20 principal components were selected for linear discriminant analysis (LDA). The number of PCs was determined by selecting the model with the lowest area under the ROC curve (AUC) (Supp Fig.S8). Classification of samples into population using PCA-LDA (C) and random forest models (D), showing percentage of samples classified to each group, standard error of the mean (SEM) and the percentage range across all replicates. Models were built and tested 20 times each with a different set of training (70%) and test (30%) data. Accuracy percentages, SEM and range were averaged across all 20 replicates. The LDA-PCA classification model had a higher accuracy (87% ± 0.02) than the random forest model (82% ± 0.02), correctly assigning 87% of individuals to their respective resistance status. Random forest models were built using 10 PCs to obtain the highest accuracy of models tested (Supp Fig.S9).

**Fig. 6:**
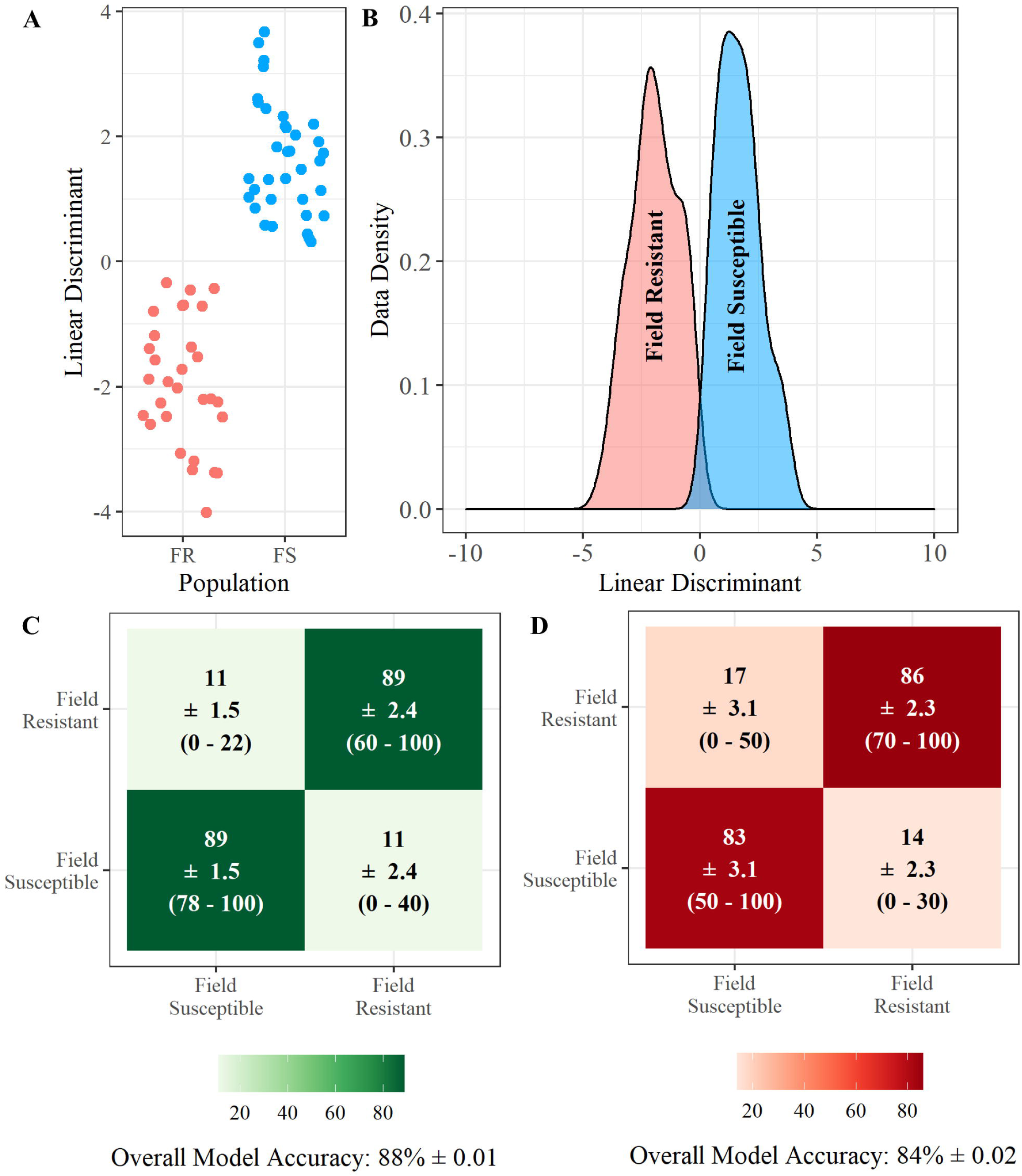
REIMS discrimination of field resistant and field susceptible *Ae. aegypti* larvae. Combined PCA-LDA separation of the resistant and field susceptible populations using REIMS mass spectra (A & B). Dimension reduction was conducted using principal components analysis (PCA), 20 principal components were selected for linear discriminant analysis (LDA). The number of PCs was determined by selecting the model with the lowest area under the ROC curve (AUC) (Supp Fig.S10). Classification of samples into population using PCA-LDA (C) and random forest models (D), showing percentage of samples classified to each group, standard error of the mean (SEM) and the percentage range across all replicates. Models were built and tested 20 times each with a different set of training (70%) and test (30%) data. Accuracy percentages, SEM and range were averaged across all 20 replicates. The LDA-PCA classification model had a higher accuracy (88% ± 0.01) than the random forest model (84% ± 0.02), correctly assigning 88% of individuals to their respective resistance status. Random forest models were built using 20 PCs to obtain the highest accuracy of models tested (Supp Fig.S11).

## Discussion

Early detection of resistance in mosquito populations is key to effective IRM and in reducing its effect on transmission of disease (Dusfour et al. 2019). The current principal methods for monitoring resistance are bioassays, biochemical assays, and molecular testing. Biochemical assays and molecular testing can be used to identify resistance in mosquitoes and are also important for the identification of mechanisms conferring resistance which can be useful when deciding on the most effective control method and in the development of novel control strategies (Brogdon 1989, World Health Organization (WHO) 1998, Corbel and N’Guessan 2013, Hemingway et al. 2013, Faucon et al. 2017, Dusfour et al. 2019). Current understanding of resistance has been developed through molecular and biochemical studies which have identified common resistance mechanisms including target site insensitivity and metabolic detoxification (Hemingway et al. 2004). Identification of these resistance mechanisms has been vital to increasing understanding of resistance.

Biochemical and molecular assays are important for increasing understanding of resistance mechanisms however there is an operational need for scalable rapid identification tools which are less labour intensive thereby yielding faster results which therefore have the potential to have more direct impact on decision making in the field. Insecticide bioassays are currently the only method for phenotyping resistance in mosquitoes (World Health Organization 2013, World Health Organization (WHO) 2016). They are limited to detecting high levels of resistance only which is often too late for alternative control methods to be deployed and high level of variation between experiments is often observed (Owusu et al. 2017). Bioassays also require large numbers of mosquitoes, the availability of a comparable susceptible strain and insectary facilities (World Health Organization 2013, World Health Organization (WHO) 2016).

This study presents proof of concept for the use of rapid evaporative ionisation mass spectrometry (REIMS) as a faster tool for monitoring of insecticide resistance which has the potential to directly inform vector control decision making. The data obtained by REIMS analysis was able to categorise resistance with 85% (± 0.01) accuracy. This method also benefits from requiring no sample preparation, and rapid data acquisition. For this study relatively small sample numbers were used, but high accuracy was still obtained. Accuracy of classification models has potential to increase as the size of the training data set is increased, therefore subsequent testing with higher sample numbers may yield an even greater accuracy, however higher variability of samples (diet, ages, environmental factors etc.) would need to be included in order to produce a robust model capable of dealing with fully wild samples (Dobbin et al. 2008, Figueroa et al. 2012, Hanberry et al. 2012, Beleites et al. 2013, Luan et al. 2020). The tool was also able to differentiate between different mosquito populations with 82% (± 0.01) accuracy, suggesting other applications for the tool aside from resistance monitoring.

We also compared two different classification model types, linear discriminant analysis (LDA) and random forest (RF) both of which are commonly applied to classification of samples using REIMS data (Cameron et al. 2016, St John et al. 2017, Davidson et al. 2019, Gredell et al. 2019, Wagner et al. 2020, Sarsby et al. 2021). LDA is often the classification method of choice for spectrometry-based phenotyping, including REIMS (Bonetti 2018, D’Hue et al. 2018, Gredell et al. 2019, Kenar et al. 2019, Liu et al. 2021, Wang et al. 2021). The results of this study showed that LDA classification models were able to achieve comparable accuracy to the more complex random forest models and in the case of our data performed better. Use of a simpler but equally accurate model is important in enabling the data analysis to be accessible to a variety of personnel working within vector control. The PCA-LDA method has previously been shown to be effective at classifying groups which show large differences in biochemical profile, however for groups with more subtle differences machine learning methods may have higher accuracy than LDA (Gromski et al. 2015, Gredell et al. 2019). The higher accuracy of the LDA model used in this study comparatively to the RF model suggests that the differences in molecular profile between the groups studied; geographical origin, population type and resistance status may be distinct. This provides further promise for the use of REIMS in insecticide resistance monitoring as larger differences in lipid signatures are easier to detect than subtle differences. The use of multiple classification models to accurately classify REIMS data has previously been shown to be important due to the high complexity of REIMS data. Dimension reduction, as conducted in this study, has also been shown to be a critical step in REIMS data analysis (Gredell et al. 2019).

Whilst the REIMS method is a fast and effective method it does have some disadvantages when compared with alternative methods. The technique is destructive, meaning that the sample cannot be used for further analysis. However, application of the technique to adult mosquitoes provides the opportunity for partial dissection (e.g. leg removal) prior to REIMS which will allow for further genetic or biochemical testing. The mass spectroscopy equipment involved in REIMS is estimated to cost around $500,000 USD (Logrono 2020), whilst costs of the initial set up of REIMS facilities are high, once equipment is available the cost per sample is low due to rapid sampling turnover. Costs are also saved elsewhere without the need for high staffing costs and insectary facilities. The speed at which samples can be analysed allows for high sample turnover which therefore reduces cost, 100 mosquito larvae could be analysed, and an answer generated in as little as 2-3 hours. In other applications including cancer diagnostic REIMS has been identified to be a more cost-effective method than other molecular techniques with costs around £1.60 per sample (Paraskevaidi et al. 2020). The REIMS method identifies differences in the lipid/metabolite profile of samples however specific molecule detection is not the objective of this method, which is designed instead to detect unique patterns in mass spectrum that enable classification (Wagner et al. 2020). Whilst we propose the use of REIMS as a potential rapid resistance identification tool with direct operational impact the technique is not intended to be used for identification of the mechanisms conferring the detected resistance.

Near-infrared spectroscopy (NIRS) is another rapid technique that has been utilised for examining invertebrates which is non-destructive and cost-effective (Johnson 2020). The high sensitivity spectrometers required for NIRS analysis cost an estimated $45,000 - $60,000 USD (Ferguson et al. 2009, Fernandes et al. 2018, Maia et al. 2019). The technique has been used successfully to differentiate mosquito species and age (Ferguson et al. 2009, Sikulu et al. 2010, 2011, Dowell et al. 2015, González Jiménez et al. 2019) and can also identify mosquitoes which are infected with arboviruses, *Plasmodium* and *Wolbachia* (Sikulu-Lord et al. 2016, Fernandes et al. 2018, Maia et al. 2019). The ability of NIRS to estimate age of mosquitoes has also been applied to the detection of insecticide resistance (Sikulu et al. 2014, Lambert et al. 2018), as insecticide resistance has been shown to decrease with age (Lines and Nassor 1991, Rajatileka et al. 2011, Jones et al. 2012). However there has been no studies which investigate the use of NIRS to directly measure insecticide resistance. The accuracy of NIRS for mosquito species determination is reported to be 78 – 90% (Ferguson et al. 2009, Sikulu et al. 2010, 2011, González Jiménez et al. 2019), lower than the 91 – 100% REIMS accuracy for species differentiation in *Drosophila* (Wagner et al. 2020). As NIRS has not been used to directly monitor insecticide resistance, comparisons between REIMS and NIRS accuracy for this purpose cannot be made.

This study focussed on identifying resistance to temephos however resistance to one insecticide rarely occurs in isolation. *Ae. aegypti* from both Cúcuta and Bello have previously been reported to have resistance to the pyrethroid permethrin and Cúcuta also to lambda-cyhalothrin (Granada et al. 2021). Whilst the current study provides proof of concept for the potential use of REIMS in identifying resistance, further study is needed to establish whether the tool can be used to differentiate between resistance to different insecticides, an application which could be beneficial to vector control programmes. Knock down resistance (*kdr)*, mutations in the sodium channel gene frequently associated with pyrethroid resistance, has also been reported in *Ae. aegypti* from Bello and Cúcuta. The varying frequencies of *kdr* alleles demonstrates that these populations are not genetically homogenous (Granada et al. 2021). Whilst gaining an understanding of the genetic basis of resistance is important (e.g. in tracking resistance and development of new interventions) it has little direct impact on the rapid decision making needed in the field (Vontas and Mavridis 2019). This study aims to provide a method which fulfils the need for more rapid resistance phenotyping tools to contribute to existing strategies without delving into the mechanisms contributing to this however there is also a further potential application of REIMS in investigating the genetic basis of resistance.

To reduce the confounding effects of phenotypic differences between populations unrelated to resistance, this study used two different susceptible populations of *Ae. aegypti*, one of field origin and a lab strain. Whilst this experimental design does reduce these confounding effects, as shown when comparing gene expression (Morgan et al. 2021), it cannot mitigate them completely and therefore other phenotypic differences between populations may be contributing to the high REIMS accuracy. This cannot be fully avoided when using field collected populations of mosquitoes.

Further testing is required to establish sensitivity of REIMS to more granular levels of resistance, resistance in other medically important mosquito species, resistance to a variety of insecticides as well as resistance in adult mosquitoes. Determining whether the preservation method of mosquito samples (e.g., desiccation, storage temperatures, fixation) affects results also has implications for field application. The results presented here identified REIMS as a promising alternative tool for the identification of insecticide resistance in mosquitoes. REIMS and similar modern phenotyping methods should be standardised and incorporated into existing insecticide resistance management strategies.

## Supplementary Material

**Supplementary File 1: R Code for analysing REIMS data**. R coding for analysing REIMS data matrices, following data binning in OMB, using LDA and random forest classification models.

**Supplementary Table 1: The REIMS data matrices**. REIMS data following binning in OMB. Data organised by population type, population, and resistance status. Mass spectra displayed in 0.1 *m/z* wide bins from 50 – 1200 *m/z*.

**Supplementary Figures S1 – S11: Supplementary figures and figure legends**. Separation of data with random group assignment (Fig S1). LDA and RF validation plots (Fig S2-S11).

## Acknowledgments

The authors thank The University of Liverpool for the support to JES-S, IW and RJB, Edge Hill University for support to JM and CS and University of Antioquia for support to OT-C. The REIMS instrumentation was supported by a grant from the Biological and Biotechnological Sciences Research Council (BB/L014793/1) to RJB.

## Author Contributions

JM: data curation; formal analysis; investigation; methodology; resources; visualization; writing – original draft, writing – review & editing. JES-S: conceptualization; supervision; writing – review and editing. IW: methodology, investigation, writing – review and editing. RJB: methodology; resources; writing – review and editing. OT-C: resources; writing – review and editing. CS: conceptualization; supervision; writing – review and editing.

---

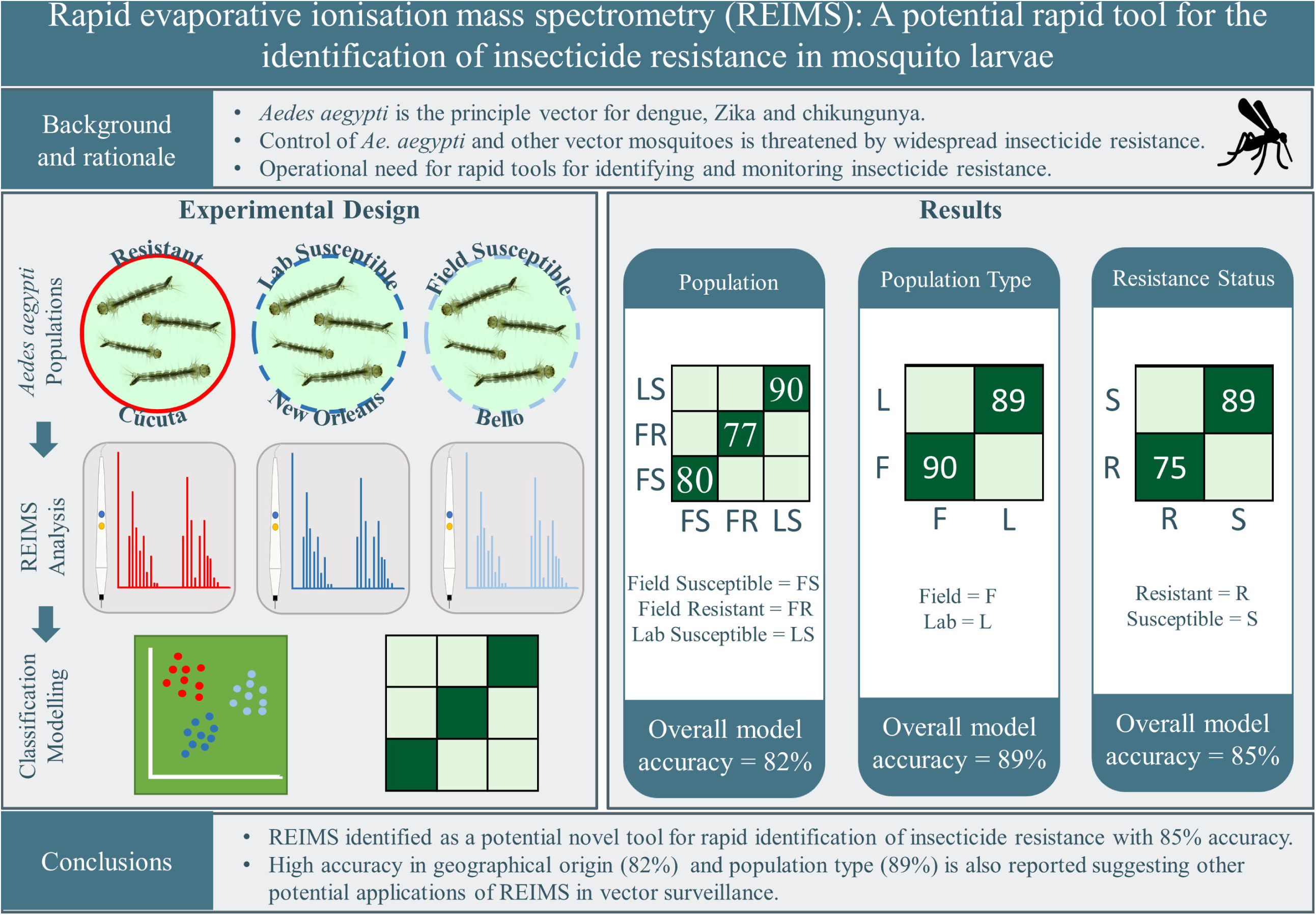

