## Supplementary Fig. for "Rapid evaporative ionisation mass spectrometry (REIMS): A potential and rapid tool for the identification of insecticide resistance in mosquito larvae"

### **Supplementary Figures**

#### **Supplementary Fig S1**

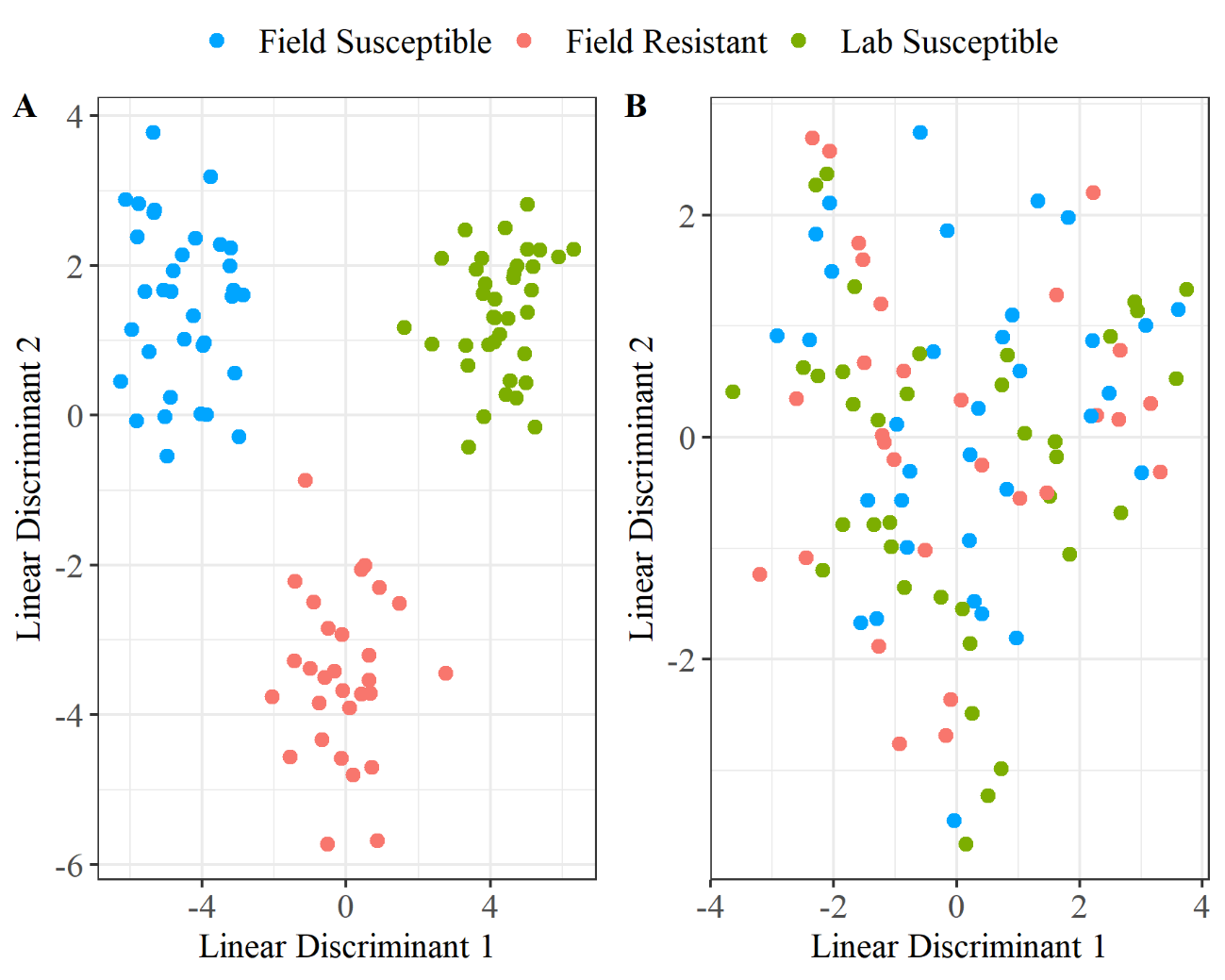

**Supplementary Fig S1:** PCA-LDA showed clear separation of groups when correct classifications were used (A). When randomly assigned classifications were used (B) no separation could be observed. This demonstrates that the observed separation of correctly assigned classifications is due to variation between populations and not due to chance.

**Supplementary Fig S2**

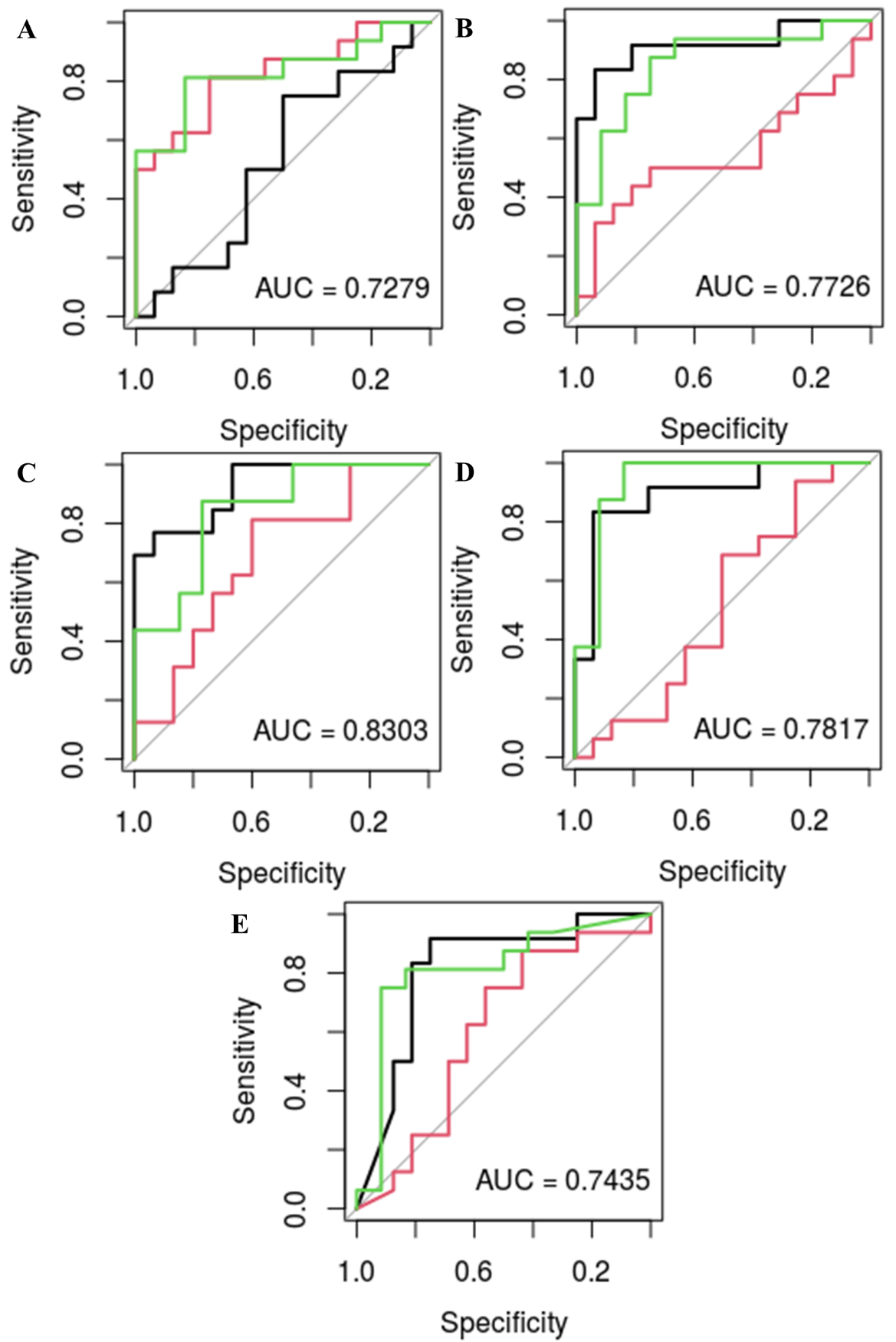

**Supplementary Fig S2: Validation of LDA built using geographical origins.** Receiver operating characteristic (ROC) plots including area under ROC curve (AUC) for LDA models built using geographical origin of samples with varying levels of principle components selected by PCA. LDA models built with 10 PCs (A), 20 PCs (B), 40 PCs (C), 60 PCs (D) and 80 PCs (E). The model with the highest AUC score was the model built using 40 principle components (C) therefore 40 PCs were used for LDA classification of data by geographical origin, the results of which are detailed in Fig 2A &B.

#### **Supplementary Fig S3**

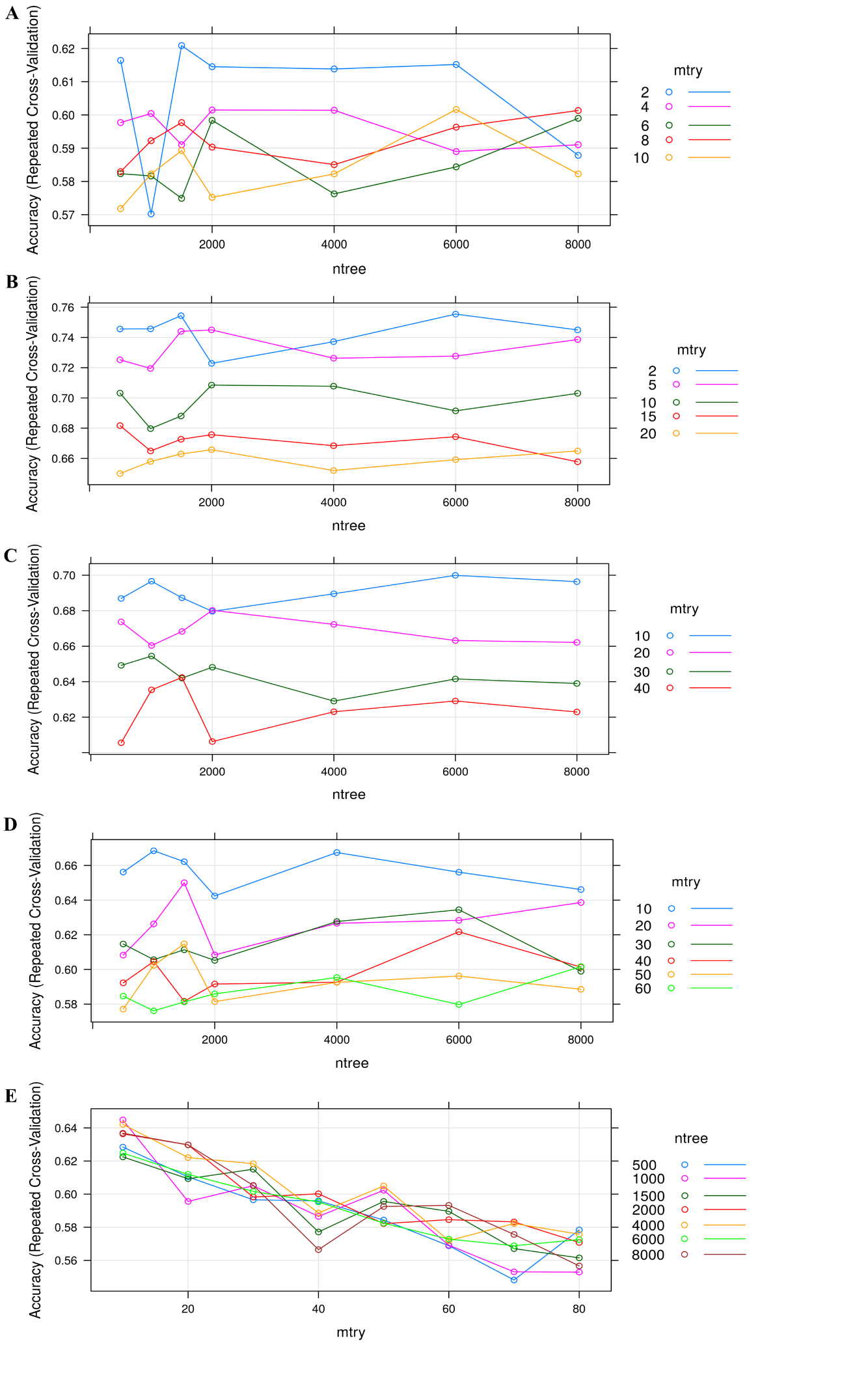

**Supplementary Fig S3: Validation of random forest built using geographical origins.** Model accuracy plots for random forest models built using geographical origin of samples with varying levels of principle components selected by PCA varying numbers of variables available for splitting at each tree node (mtry) and various tree numbers. Random forest models built with 10 PCs (A), 20 PCs (B), 40 PCs (C), 60 PCs (D) and 80 PCs (E). The model with the highest accuracy was the model built using 20 principle components (C) with mtry = 2 and ntree = 6000 therefore 20 PCs were used for random model classification of data by geographical origin using mtry = 2 and ntree = 6000 the results of which are detailed in Fig 2C.

#### **Supplementary Fig S4**

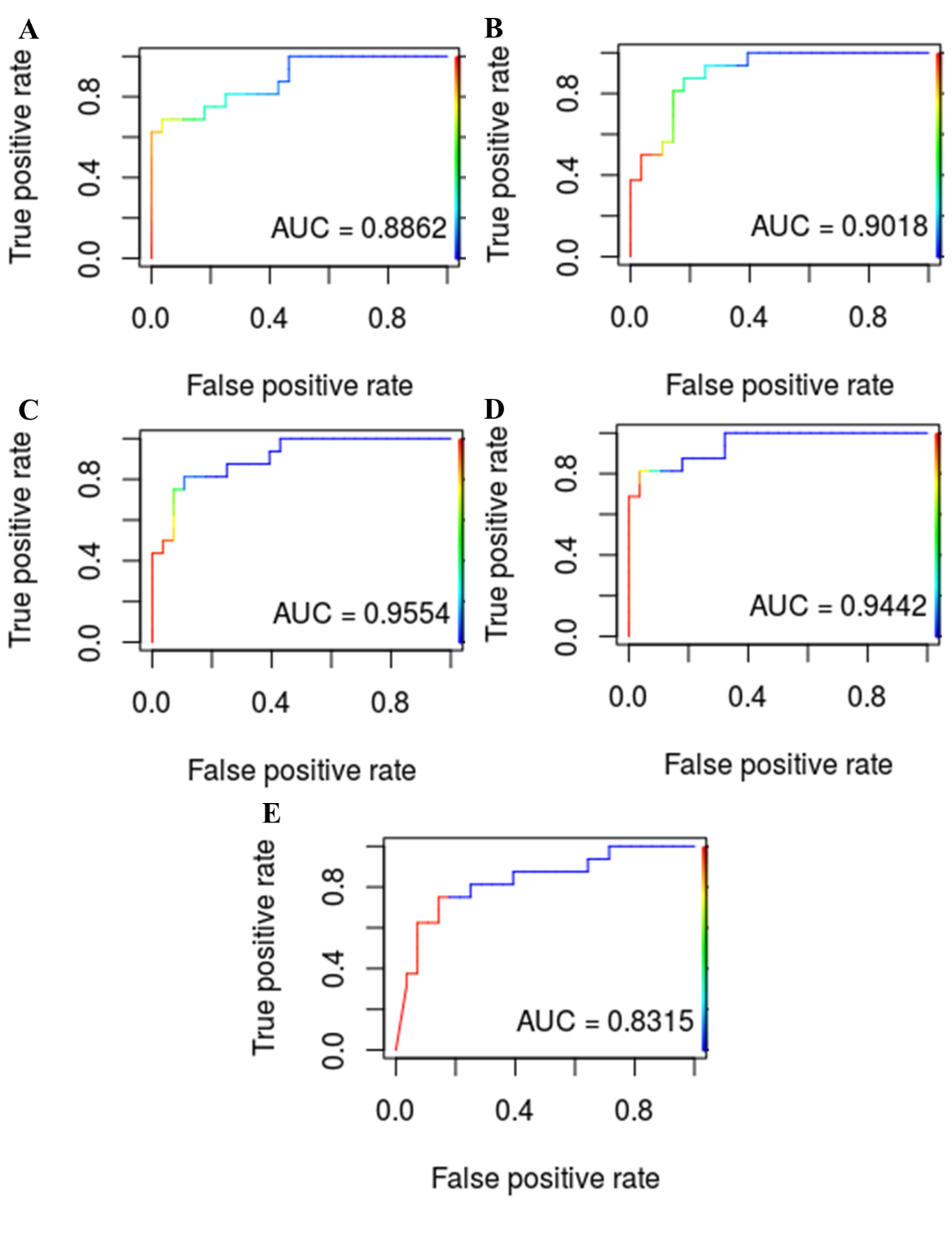

**Supplementary Fig S4: Validation of LDA built using population type.** Receiver operating characteristic (ROC) plots including area under ROC curve (AUC) for LDA models built using population type (lab or field) of samples with varying levels of principle components selected by PCA. LDA models built with 10 PCs (A), 20 PCs (B), 40 PCs (C), 60 PCs (D) and 80 PCs (E). The model with the highest AUC score was the model built using 40 principle components (C) therefore 40 PCs were used for LDA classification of data by population type, the results of which are detailed in Fig 3A &B.

**Supplementary Fig S5**

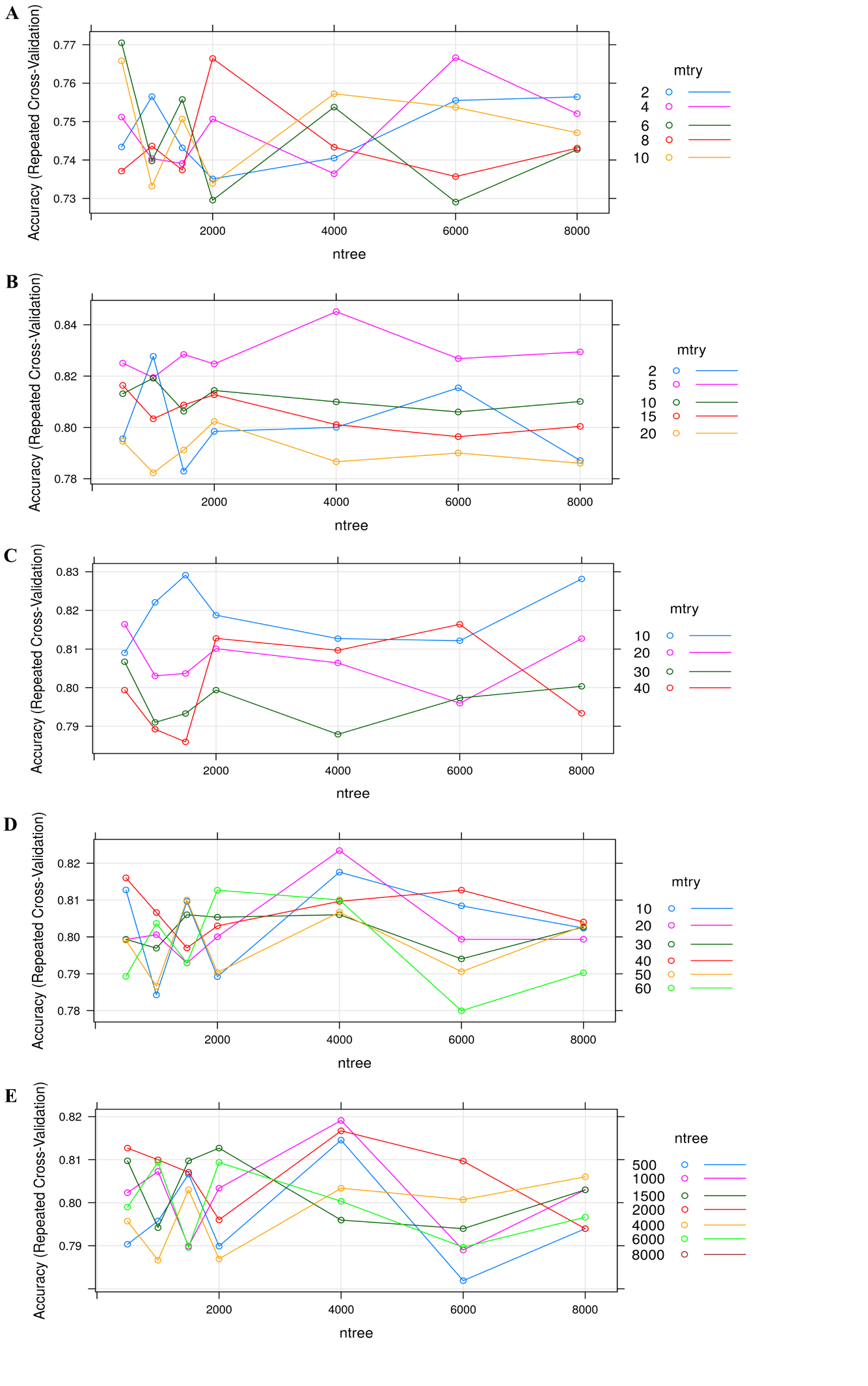

**Supplementary Fig S5: Validation of random forest built using population type.** Model accuracy plots for random forest models built using population type (lab or field) of samples with varying levels of principle components selected by PCA varying numbers of variables available for splitting at each tree node (mtry) and various tree numbers. Random forest models built with 10 PCs (A), 20 PCs (B), 40 PCs (C), 60 PCs (D) and 80 PCs (E). The model with the highest accuracy was the model built using 20 principle components (C) with mtry = 5 and ntree = 4000 therefore 20 PCs were used for random model classification of data by geographical origin using mtry = 5 and ntree = 4000 the results of which are detailed in Fig 3C.

#### **Supplementary Fig S6**

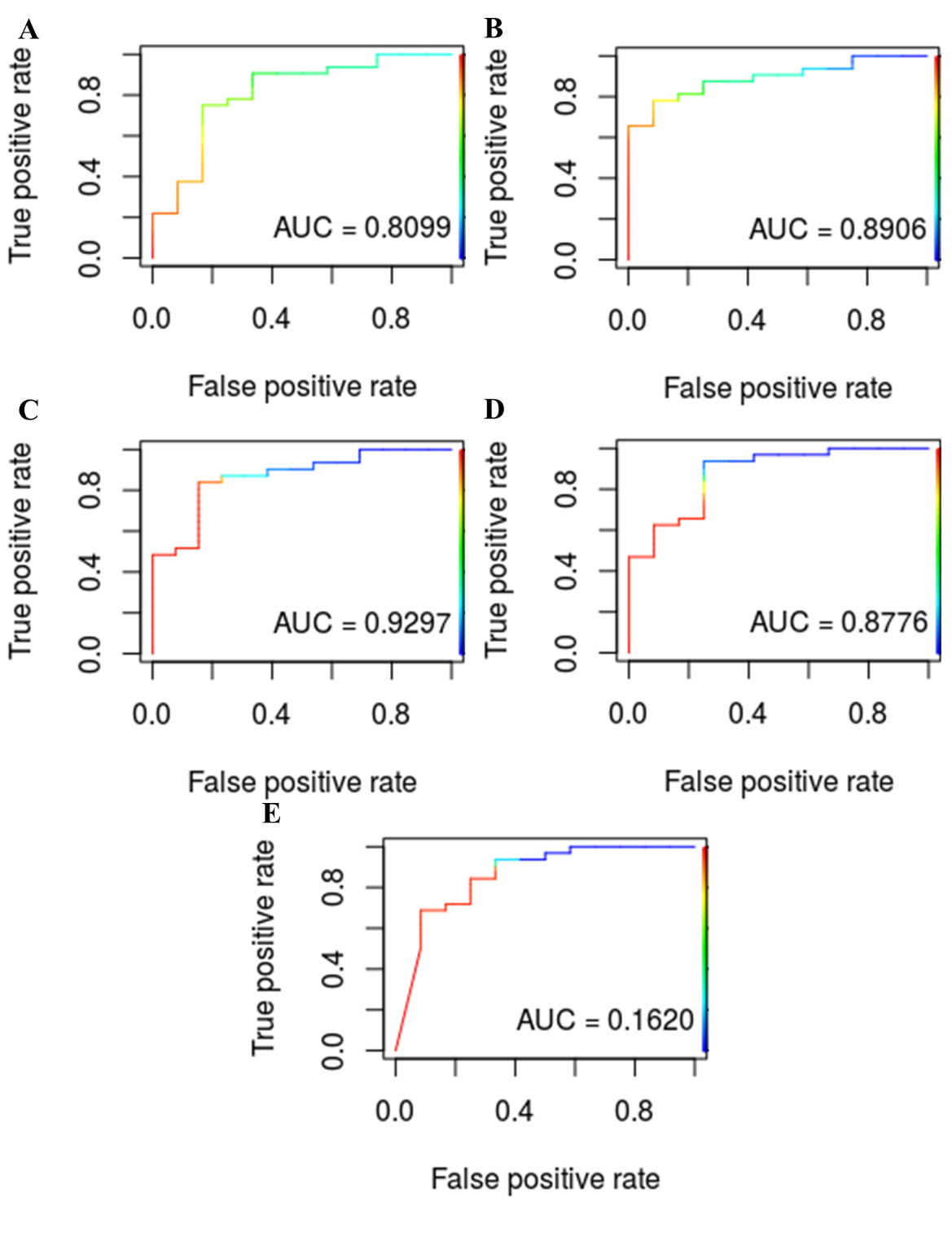

**Supplementary Fig S6: Validation of LDA built using insecticide resistance status.** Receiver operating characteristic (ROC) plots including area under ROC curve (AUC) for LDA models built using insecticide resistance status of samples with varying levels of principle components selected by PCA. LDA models built with 10 PCs (A), 20 PCs (B), 40 PCs (C), 60 PCs (D) and 80 PCs (E). The model with the highest AUC score was the model built using 40 principle components (C) therefore 40 PCs were used for LDA classification of data by insecticide resistance status, the results of which are detailed in Fig 4A &B.

#### **Supplementary Fig S7**

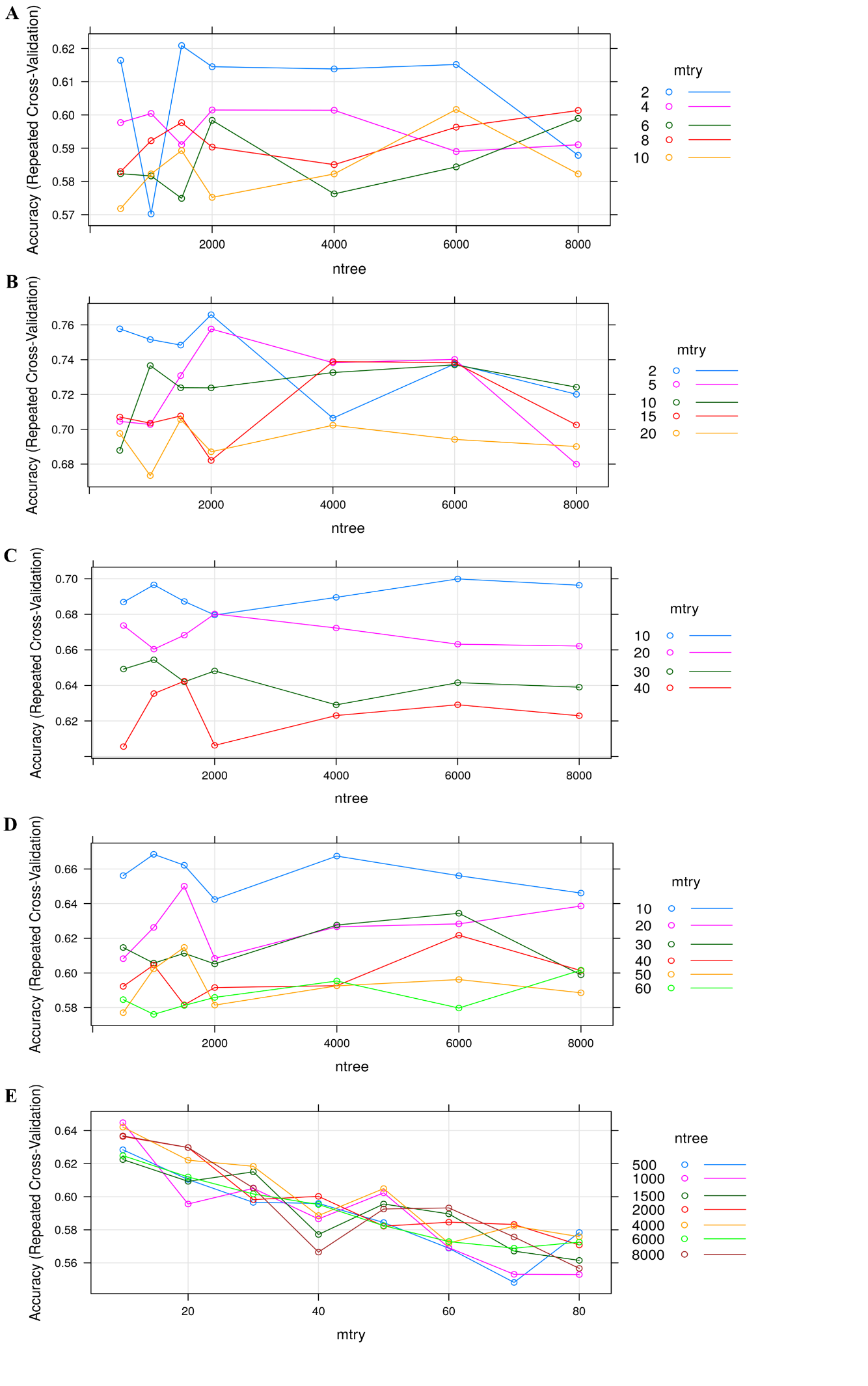

**Supplementary Fig S7: Validation of random forest built using insecticide resistance status.** Model accuracy plots for random forest models built using insecticide resistance status of samples with varying levels of principle components selected by PCA varying numbers of variables available for splitting at each tree node (mtry) and various tree numbers. Random forest models built with 10 PCs (A), 20 PCs (B), 40 PCs (C), 60 PCs (D) and 80 PCs (E). The model with the highest accuracy was the model built using 20 principle components (C) with mtry = 2 and ntree = 2000 therefore 20 PCs were used for random model classification of data by geographical origin using mtry = 2 and ntree = 2000 the results of which are detailed in Fig 4C.

#### Supplementary Fig S8

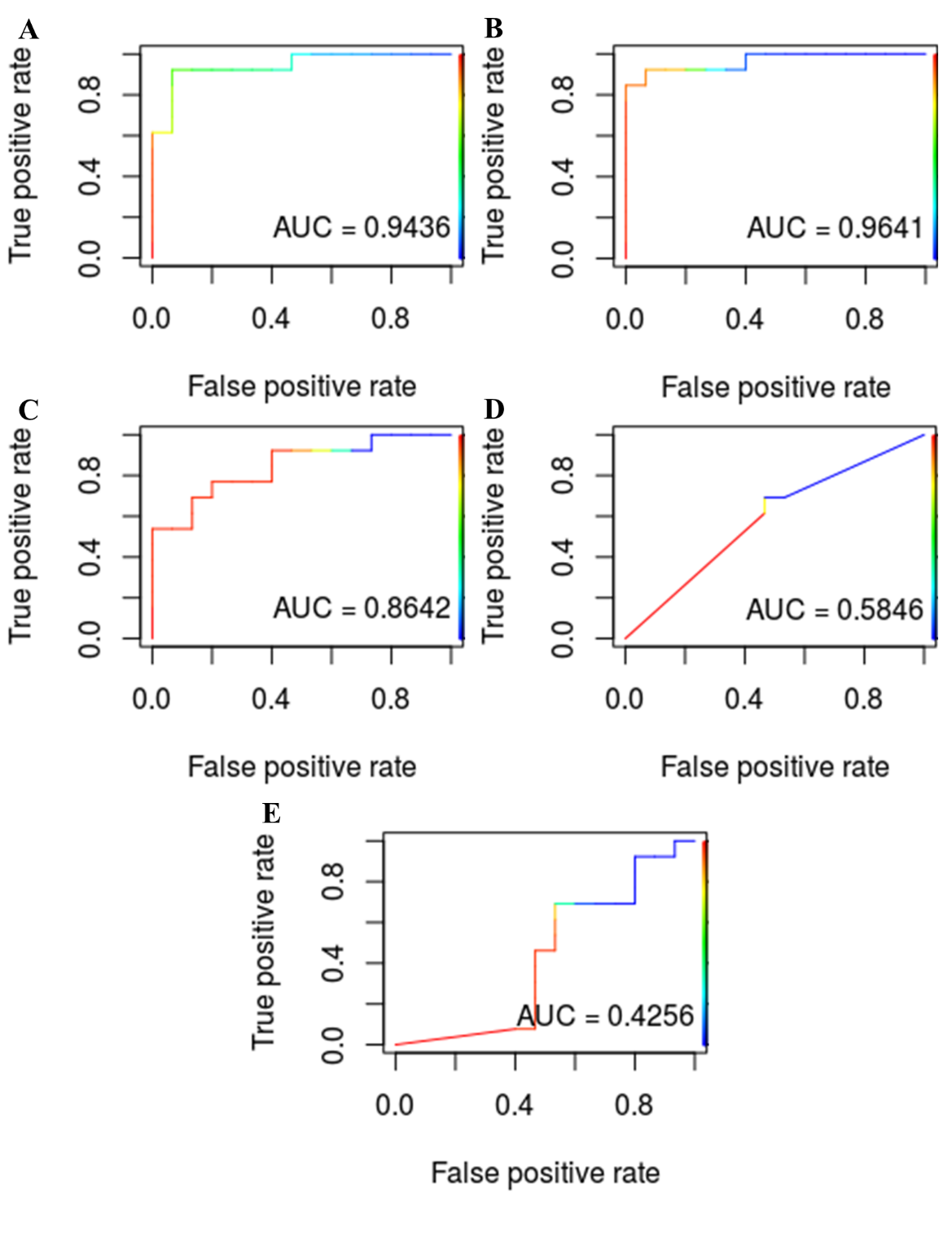

**Supplementary Fig S8: Validation of LDA built using insecticide resistance status with field resistant and lab susceptible groups only.** Receiver operating characteristic (ROC) plots including area under ROC curve (AUC) for LDA models built using insecticide resistance status of samples (FR and LS) with varying levels of principle components selected by PCA. LDA models built with 10 PCs (A), 20 PCs (B), 40 PCs (C), 60 PCs (D) and 80 PCs (E). The model with the highest AUC score was the model built using 20 principle components (C) therefore 20 PCs were used for LDA classification of data by insecticide resistance status, the results of which are detailed in Fig 5A &B.

#### **Supplementary Fig S9**

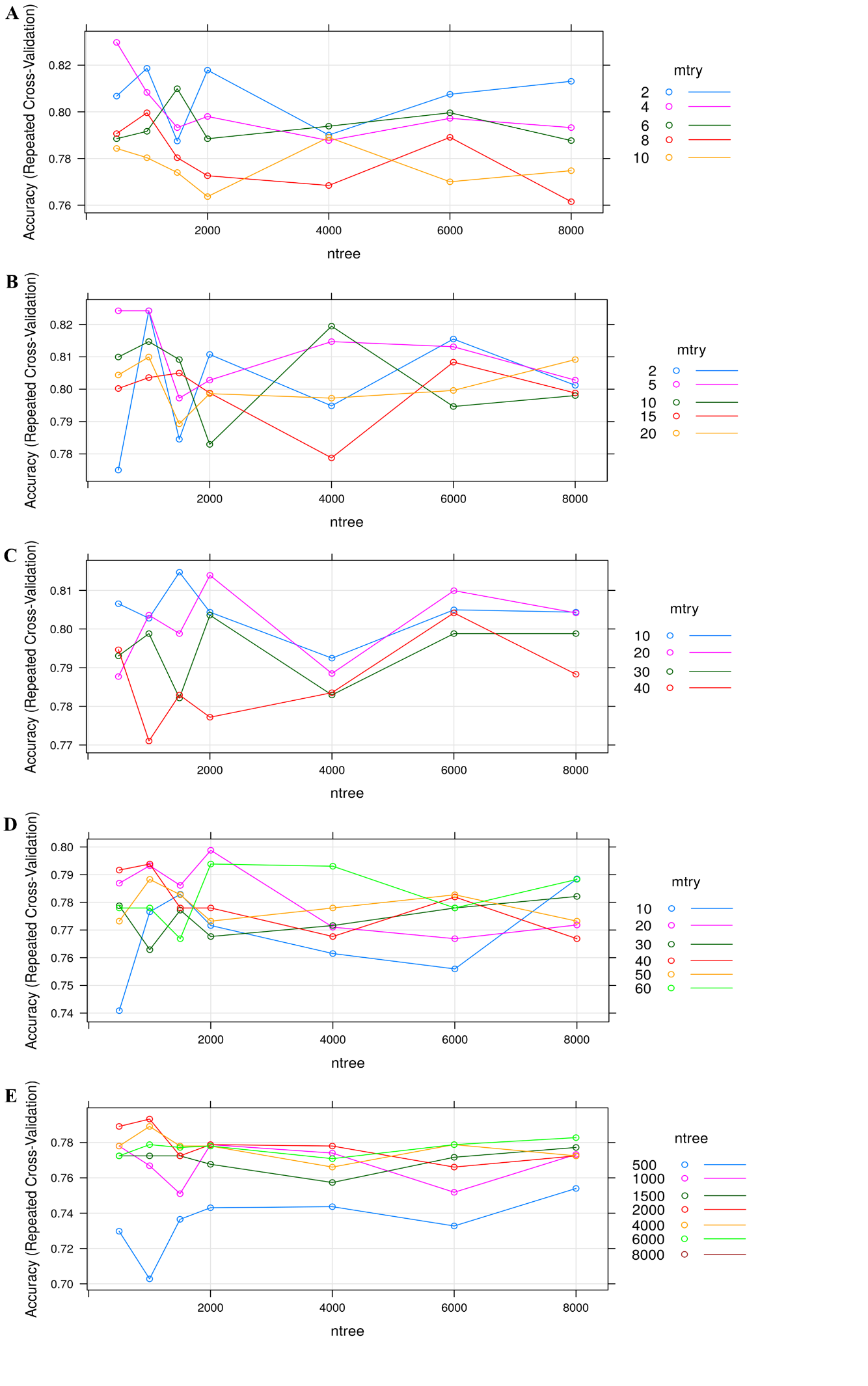

**Supplementary Fig S9: Validation of random forest built using insecticide resistance status with field resistant and lab susceptible groups only.** Model accuracy plots for random forest models built using insecticide resistance status of samples (FR and LS) with varying levels of principle components selected by PCA varying numbers of variables available for splitting at each tree node (mtry) and various tree numbers. Random forest models built with 10 PCs (A), 20 PCs (B), 40 PCs (C), 60 PCs (D) and 80 PCs (E). The model with the highest accuracy was the model built using 10 principle components (C) with mtry = 4 and ntree = 500 therefore 10 PCs were used for random model classification of data by geographical origin using mtry = 4 and ntree = 500 the results of which are detailed in Fig 5C.

#### **Supplementary Fig S10**

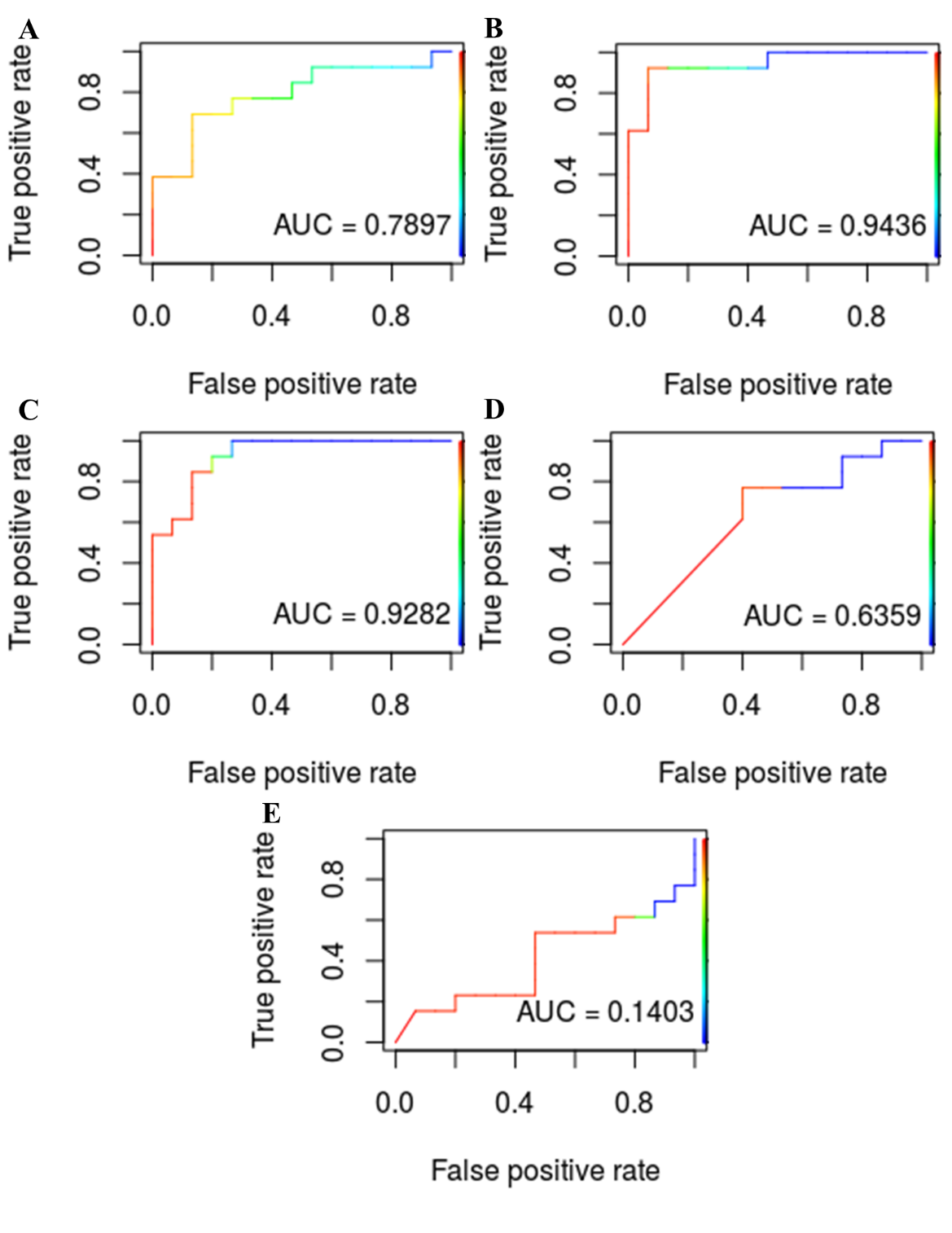

**Supplementary Fig S10: Validation of LDA built using insecticide resistance status with field resistant and field susceptible groups only.** Receiver operating characteristic (ROC) plots including area under ROC curve (AUC) for LDA models built using insecticide resistance status of samples (FR and FS) with varying levels of principle components selected by PCA. LDA models built with 10 PCs (A), 20 PCs (B), 40 PCs (C), 60 PCs (D) and 80 PCs (E). The model with the highest AUC score was the model built using 20 principle components (C) therefore 20 PCs were used for LDA classification of data by insecticide resistance status, the results of which are detailed in Fig 6A &B.

#### **Supplementary Fig S11**

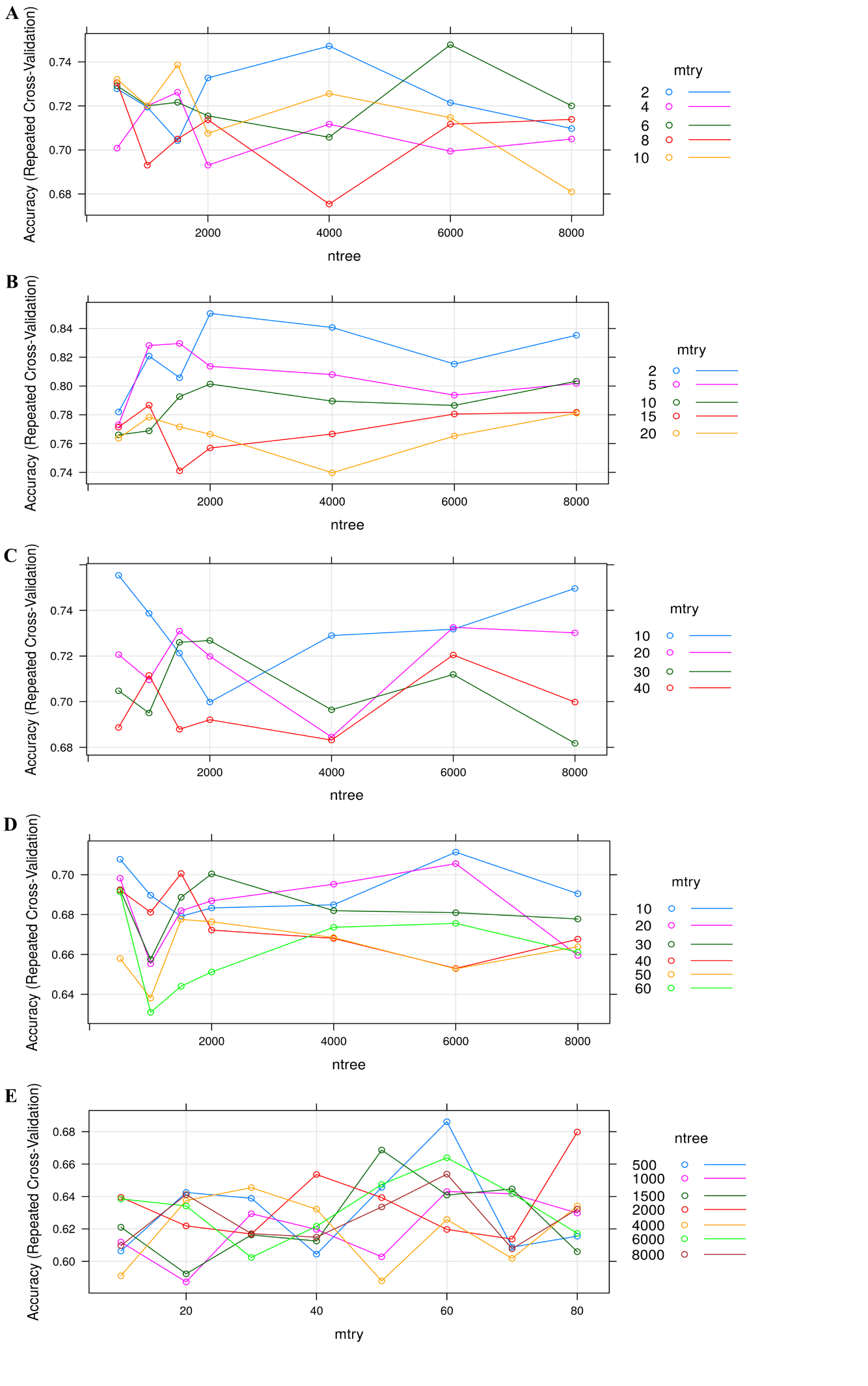

**Supplementary Fig S11: Validation of random forest built using insecticide resistance status with field resistant and field susceptible groups only.** Model accuracy plots for random forest models built using insecticide resistance status of samples (FR and FS) with varying levels of principle components selected by PCA varying numbers of variables available for splitting at each tree node (mtry) and various tree numbers. Random forest models built with 10 PCs (A), 20 PCs (B), 40 PCs (C), 60 PCs (D) and 80 PCs (E). The model with the highest accuracy was the model built using 20 principle components (C) with mtry = 2 and ntree = 2000 therefore 20 PCs were used for random model classification of data by geographical origin using mtry = 2 and ntree = 2000 the results of which are detailed in Fig 6C.
