## Supplementary File 1 for "Rapid evaporative ionisation mass spectrometry (REIMS): A potential and rapid tool for the identification of insecticide resistance in mosquito larvae"

####Reproducable script for analysing REIMS data (Morgan et al. 2022).
###The method here uses PCA for dimensionality reduction followed by LDA and random forest classification models.
###Plots of the LDA are also produced.

##Install and load required packages.
install.packages ("factoextra")
install.packages ("MASS")
install.packages ("ggplot2")
install.packages ("ggpubr")
install.packages ("randomForest")
install.packages ("caret")
install.packages ("xlsx")
install.packages("ROCR")
install.packages ("pROC")

library ("factoextra")
library ("MASS")
library ("ggplot2")
library ("ggpubr")
library ("randomForest")
library ("caret")
library ("xlsx")
library ("ROCR")
library ("pROC")

##Import the dataset.
Data <- read.table("PATH_TO_DATASET/Data.txt", header = TRUE, check.names = FALSE)

##If the explanatory variable (group) is not numeric it must be converted to numeric. 
#This will give each category a number, make not of which category is assigned which number as it will be needed later.
Data$Group = as.numeric(as.factor(Data$Group))
Group <- Data$Group

#convert data to data frame.
Data <- as.data.frame(Data)

##Conduct PCA in order to reduce the number of dimensions that will be used for LDA and RF.
#Run PCA, view summary of the results.
PCA <- prcomp (Data)
summary (PCA)

##The PCA can be explored by plotting scree plots.
#Scree Plot for % explained variance
fviz_eig(PCA, addlabels = TRUE)

#Scree Plot for Eigenvalues.
fviz_eig(PCA, choice = "eigenvalue",addlabels = TRUE)

##Use the PCA to reduce dimensions
#The total number of dimensions will depend on the number of observations in the dataset.
#Try reducing to different numbers of dimensions by adjusting the numbers in the square brackets, having a variety of different numbers of PCs will be helpful when building optimal models later. 
#In this example dimensions will be reduced to 10,20,40,60,80,100.

#10 dimensions.
Data_10_DR <- PCA$x[,1:10]
head (Data_10_DR)

#20 dimensions.
Data_20_DR <- PCA$x[,1:20]
head (Data_20_DR)

#40 dimensions.
Data_40_DR <- PCA$x[,1:40]
head (Data_40_DR)

#60 dimensions.
Data_60_DR <- PCA$x[,1:60]
head (Data_60_DR)

#80 dimensions.
Data_80_DR <- PCA$x[,1:80]
head (Data_80_DR)

#100 dimensions.
Data_100_DR <- PCA$x[,1:100]
head (Data_100_DR)

##Add the group information to the dimension reduced datasets.
#Ensure that the number in square brackets in the colname command is one more than the total number of dimensions.

#10 dimensions.
Data_10_DR <- cbind(Data_10_DR, as.numeric(Data$Group))
colnames(Data_10_DR) [11] <- "Group"
head (Data_10_DR)

#20 dimensions.
Data_20_DR <- cbind(Data_20_DR, as.numeric(Data$Group))
colnames(Data_20_DR) [21] <- "Group"
head (Data_20_DR)

#40 dimensions.
Data_40_DR <- cbind(Data_40_DR, as.numeric(Data$Group))
colnames(Data_40_DR) [41] <- "Group"
head (Data_40_DR)

#60 dimensions.
Data_60_DR <- cbind(Data_60_DR, as.numeric(Data$Group))
colnames(Data_60_DR) [61] <- "Group"
head (Data_60_DR)

#80 dimensions.
Data_80_DR <- cbind(Data_80_DR, as.numeric(Data$Group))
colnames(Data_80_DR) [81] <- "Group"
head (Data_80_DR)

#100 dimensions.
Data_100_DR <- cbind(Data_100_DR, as.numeric(Data$Group))
colnames(Data_100_DR) [101] <- "Group"
head (Data_100_DR)

#Convert the reduced data sets into data frames and make group a factor.

#10 dimensions.
Data_10_DR <- as.data.frame(Data_10_DR)
Data_10_DR$Group <- as.factor(Data_10_DR$Group)

#20 dimensions.
Data_20_DR <- as.data.frame(Data_20_DR)
Data_20_DR$Group <- as.factor(Data_20_DR$Group)

#40 dimensions.
Data_40_DR <- as.data.frame(Data_40_DR)
Data_40_DR$Group <- as.factor(Data_40_DR$Group)

#60 dimensions.
Data_60_DR <- as.data.frame(Data_60_DR)
Data_60_DR$Group <- as.factor(Data_60_DR$Group)

#80 dimensions.
Data_80_DR <- as.data.frame(Data_80_DR)
Data_80_DR$Group <- as.factor(Data_80_DR$Group)

#100 dimensions.
Data_100_DR <- as.data.frame(Data_100_DR)
Data_100_DR$Group <- as.factor(Data_100_DR$Group)

##LDA is going to be used as a classifier.
##For classification models the data will be split randomly into 70% training and 30% test data. These proportions can be adjusted to suit data.
##The LDA should be repeated at least 10x with different train and test data.

##Use the first set of train/test data determine optimal number of PCs to use in LDA.This should be completed for the data sets with differing numbers of principle comonents compared above.

#Testing 10 PCs
#Define train and test split (70% Train/30% Test) - randomly.
set.seed(11) #This is important to insure reproducibility of results, changing this number also allows for different random splits of data.
Data_10_train_size <- floor(0.7 * nrow(Data_10_DR)) #To adjust proportion of data to be assigned to training change the value (0.7) here.
set.seed(11)
Data_10_train_ind <- sample(nrow(Data_10_DR), size = Data_10_train_size)
set.seed(11)
Data_10_train_data <- as.data.frame(Data_10_DR[Data_10_train_ind,])
set.seed(11)
Data_10_test_data <- as.data.frame(Data_10_DR[-Data_10_train_ind,])

#Conduct linear discriminant analysis
set.seed (11)
Data_10_LDA <- lda(Group ~ ., data = Data_10_train_data)

#Use the LDA to make predictions
set.seed (11)
Data_10_LDA_predict <- predict(Data_10_LDA, newdata = Data_10_test_data, type = "prob")
head(Data_10_LDA_predict)

#Create and view the confusion matrix.
set.seed (11)
Data_table_1_10= table (Data_10_LDA_predict$class,Data_10_test_data$Group)
Data_table_1_10 <- confusionMatrix(Data_table_1_10)
Data_table_1_10

#Create ROC curve. Use this code when data has >2 different groups. 
Data_ROC_10 <- multiclass.roc (Data_10_test_data$Group, Data_10_LDA_predict$posterior [,2])
rs_10 <- Data_ROC_10[['rocs']]
Data_ROC_10_plot <- plot.roc(rs_10[[1]])
sapply(2:length (rs_10), function (i) lines.roc(rs_10[[i]], col = i))

#Create ROC curve. Use this code when data has only 2 different groups. 
pred_Data_10 <- prediction(Data_10_LDA_predict$posterior[,2], Data_10_test_data$Group) 
perf_Data_10 <- performance(pred_Data_10,"tpr","fpr")
plot(perf_Data_10,colorize=TRUE)

#Calculate AUC score, this will be used to determine the most effective model
auc (Data_ROC_10)
text("AUC = INSERTAUCSCOREHERE",  x = 0.3, y = 0.1) #The AUC score can be added to the ROC plot, adjust its position by altering the x and y values here.

#This plot should be saved for future reference. 

#Testing 20 PCs
#Define train and test split (70% Train/30% Test) - randomly.
set.seed(11) #This is important to insure reproducibility of results, changing this number also allows for different random splits of data.
Data_20_train_size <- floor(0.7 * nrow(Data_20_DR)) #To adjust proportion of data to be assigned to training change the value (0.7) here.
set.seed(11)
Data_20_train_ind <- sample(nrow(Data_20_DR), size = Data_20_train_size)
set.seed(11)
Data_20_train_data <- as.data.frame(Data_20_DR[Data_20_train_ind,])
set.seed(11)
Data_20_test_data <- as.data.frame(Data_20_DR[-Data_20_train_ind,])

#Conduct linear discriminant analysis
set.seed (11)
Data_20_LDA <- lda(Group ~ ., data = Data_20_train_data)

#Use the LDA to make predictions
set.seed (11)
Data_20_LDA_predict <- predict(Data_20_LDA, newdata = Data_20_test_data, type = "prob")
head(Data_20_LDA_predict)

#Create and view the confusion matrix.
set.seed (11)
Data_table_1_20= table (Data_20_LDA_predict$class,Data_20_test_data$Group)
Data_table_1_20 <- confusionMatrix(Data_table_1_20)
Data_table_1_20

#Create ROC curve. Use this code when data has >2 different groups. 
Data_ROC_20 <- multiclass.roc (Data_20_test_data$Group, Data_20_LDA_predict$posterior [,2])
rs_20 <- Data_ROC_20[['rocs']]
Data_ROC_20_plot <- plot.roc(rs_20[[1]])
sapply(2:length (rs_20), function (i) lines.roc(rs_20[[i]], col = i))

#Create ROC curve. Use this code when data has only 2 different groups. 
pred_Data_10 <- prediction(Data_10_LDA_predict$posterior[,2], Data_10_test_data$Group) 
perf_Data_10 <- performance(pred_Data_10,"tpr","fpr")
plot(perf_Data_10,colorize=TRUE)

#Calculate AUC score, this will be used to determine the most effective model
auc (Data_ROC_20)
text("AUC = INSERTAUCSCOREHERE",  x = 0.3, y = 0.1) #The AUC score can be added to the ROC plot, adjust its position by altering the x and y values here.

#ROC plot should be saved for future reference. 

#Testing 40 PCs
#Define train and test split (70% Train/30% Test) - randomly.
set.seed(11) #This is important to insure reproducibility of results, changing this number also allows for different random splits of data.
Data_40_train_size <- floor(0.7 * nrow(Data_40_DR)) #To adjust proportion of data to be assigned to training change the value (0.7) here.
set.seed(11)
Data_40_train_ind <- sample(nrow(Data_40_DR), size = Data_40_train_size)
set.seed(11)
Data_40_train_data <- as.data.frame(Data_40_DR[Data_40_train_ind,])
set.seed(11)
Data_40_test_data <- as.data.frame(Data_40_DR[-Data_40_train_ind,])

#Conduct linear discriminant analysis
set.seed (11)
Data_40_LDA <- lda(Group ~ ., data = Data_40_train_data)

#Use the LDA to make predictions
set.seed (11)
Data_40_LDA_predict <- predict(Data_40_LDA, newdata = Data_40_test_data, type = "prob")
head(Data_40_LDA_predict)

#Create and view the confusion matrix.
set.seed (11)
Data_table_1_40= table (Data_40_LDA_predict$class,Data_40_test_data$Group)
Data_table_1_40 <- confusionMatrix(Data_table_1_40)
Data_table_1_40

#Create ROC curve. Use this code when data has >2 different groups. 
Data_ROC_40 <- multiclass.roc (Data_40_test_data$Group, Data_40_LDA_predict$posterior [,2])
rs_40 <- Data_ROC_40[['rocs']]
Data_ROC_40_plot <- plot.roc(rs_40[[1]])
sapply(2:length (rs_40), function (i) lines.roc(rs_40[[i]], col = i))

#Create ROC curve. Use this code when data has only 2 different groups. 
pred_Data_10 <- prediction(Data_10_LDA_predict$posterior[,2], Data_10_test_data$Group) 
perf_Data_10 <- performance(pred_Data_10,"tpr","fpr")
plot(perf_Data_10,colorize=TRUE)

#Calculate AUC score, this will be used to determine the most effective model
auc (Data_ROC_40)
text("AUC = INSERTAUCSCOREHERE",  x = 0.3, y = 0.1) #The AUC score can be added to the ROC plot, adjust its position by altering the x and y values here.

#This plot should be saved for future reference. 

#Testing 60 PCs
#Define train and test split (70% Train/30% Test) - randomly.
set.seed(11) #This is important to insure reproducibility of results, changing this number also allows for different random splits of data.
Data_60_train_size <- floor(0.7 * nrow(Data_60_DR)) #To adjust proportion of data to be assigned to training change the value (0.7) here.
set.seed(11)
Data_60_train_ind <- sample(nrow(Data_60_DR), size = Data_60_train_size)
set.seed(11)
Data_60_train_data <- as.data.frame(Data_60_DR[Data_60_train_ind,])
set.seed(11)
Data_60_test_data <- as.data.frame(Data_60_DR[-Data_60_train_ind,])

#Conduct linear discriminant analysis
set.seed (11)
Data_60_LDA <- lda(Group ~ ., data = Data_60_train_data)

#Use the LDA to make predictions
set.seed (11)
Data_60_LDA_predict <- predict(Data_60_LDA, newdata = Data_60_test_data, type = "prob")
head(Data_60_LDA_predict)

#Create and view the confusion matrix.
set.seed (11)
Data_table_1_60= table (Data_60_LDA_predict$class,Data_60_test_data$Group)
Data_table_1_60 <- confusionMatrix(Data_table_1_60)
Data_table_1_60

#Create ROC curve. Use this code when data has >2 different groups. 
Data_ROC_60 <- multiclass.roc (Data_60_test_data$Group, Data_60_LDA_predict$posterior [,2])
rs_60 <- Data_ROC_60[['rocs']]
Data_ROC_60_plot <- plot.roc(rs_60[[1]])
sapply(2:length (rs_60), function (i) lines.roc(rs_60[[i]], col = i))

#Create ROC curve. Use this code when data has only 2 different groups. 
pred_Data_10 <- prediction(Data_10_LDA_predict$posterior[,2], Data_10_test_data$Group) 
perf_Data_10 <- performance(pred_Data_10,"tpr","fpr")
plot(perf_Data_10,colorize=TRUE)

#Calculate AUC score, this will be used to determine the most effective model
auc (Data_ROC_60)
text("AUC = INSERTAUCSCOREHERE",  x = 0.3, y = 0.1) #The AUC score can be added to the ROC plot, adjust its position by altering the x and y values here.

#This plot should be saved for future reference. 

#Testing 80 PCs
#Define train and test split (70% Train/30% Test) - randomly.
set.seed(11) #This is important to insure reproducibility of results, changing this number also allows for different random splits of data.
Data_80_train_size <- floor(0.7 * nrow(Data_80_DR)) #To adjust proportion of data to be assigned to training change the value (0.7) here.
set.seed(11)
Data_80_train_ind <- sample(nrow(Data_80_DR), size = Data_80_train_size)
set.seed(11)
Data_80_train_data <- as.data.frame(Data_80_DR[Data_80_train_ind,])
set.seed(11)
Data_80_test_data <- as.data.frame(Data_80_DR[-Data_80_train_ind,])

#Conduct linear discriminant analysis
set.seed (11)
Data_80_LDA <- lda(Group ~ ., data = Data_80_train_data)

#Use the LDA to make predictions
set.seed (11)
Data_80_LDA_predict <- predict(Data_80_LDA, newdata = Data_80_test_data, type = "prob")
head(Data_80_LDA_predict)

#Create and view the confusion matrix.
set.seed (11)
Data_table_1_80= table (Data_80_LDA_predict$class,Data_80_test_data$Group)
Data_table_1_80 <- confusionMatrix(Data_table_1_80)
Data_table_1_80

#Create ROC curve. Use this code when data has >2 different groups. 
Data_ROC_80 <- multiclass.roc (Data_80_test_data$Group, Data_80_LDA_predict$posterior [,2])
rs_80 <- Data_ROC_80[['rocs']]
Data_ROC_80_plot <- plot.roc(rs_80[[1]])
sapply(2:length (rs_80), function (i) lines.roc(rs_80[[i]], col = i))

#Create ROC curve. Use this code when data has only 2 different groups. 
pred_Data_10 <- prediction(Data_10_LDA_predict$posterior[,2], Data_10_test_data$Group) 
perf_Data_10 <- performance(pred_Data_10,"tpr","fpr")
plot(perf_Data_10,colorize=TRUE)

#Calculate AUC score, this will be used to determine the most effective model
auc (Data_ROC_80)
text("AUC = INSERTAUCSCOREHERE",  x = 0.3, y = 0.1) #The AUC score can be added to the ROC plot, adjust its position by altering the x and y values here.

#This plot should be saved for future reference. 

#Testing 100 PCs
#Define train and test split (70% Train/30% Test) - randomly.
set.seed(11) #This is important to insure reproducibility of results, changing this number also allows for different random splits of data.
Data_100_train_size <- floor(0.7 * nrow(Data_100_DR)) #To adjust proportion of data to be assigned to training change the value (0.7) here.
set.seed(11)
Data_100_train_ind <- sample(nrow(Data_100_DR), size = Data_100_train_size)
set.seed(11)
Data_100_train_data <- as.data.frame(Data_100_DR[Data_100_train_ind,])
set.seed(11)
Data_100_test_data <- as.data.frame(Data_100_DR[-Data_100_train_ind,])

#Conduct linear discriminant analysis
set.seed (11)
Data_100_LDA <- lda(Group ~ ., data = Data_100_train_data)

#Use the LDA to make predictions
set.seed (11)
Data_100_LDA_predict <- predict(Data_100_LDA, newdata = Data_100_test_data, type = "prob")
head(Data_100_LDA_predict)

#Create and view the confusion matrix.
set.seed (11)
Data_table_1_100= table (Data_100_LDA_predict$class,Data_100_test_data$Group)
Data_table_1_100 <- confusionMatrix(Data_table_1_100)
Data_table_1_100

#Create ROC curve. Use this code when data has >2 different groups. 
Data_ROC_100 <- multiclass.roc (Data_100_test_data$Group, Data_100_LDA_predict$posterior [,2])
rs_100 <- Data_ROC_100[['rocs']]
Data_ROC_100_plot <- plot.roc(rs_100[[1]])
sapply(2:length (rs_100), function (i) lines.roc(rs_100[[i]], col = i))

#Create ROC curve. Use this code when data has only 2 different groups. 
pred_Data_10 <- prediction(Data_10_LDA_predict$posterior[,2], Data_10_test_data$Group) 
perf_Data_10 <- performance(pred_Data_10,"tpr","fpr")
plot(perf_Data_10,colorize=TRUE)

#Calculate AUC score, this will be used to determine the most effective model
auc (Data_ROC_100)
text("AUC = INSERTAUCSCOREHERE",  x = 0.3, y = 0.1) #The AUC score can be added to the ROC plot, adjust its position by altering the x and y values here.

#This plot should be saved for future reference. 

##Saving the statistics from the different models for future reference. 
##One method is presented here but any method for saving from R can be used. 

Data_table_10_CM <- as.matrix(Data_table_1_10)
Data_table_10_OS <- as.matrix(Data_table_1_10, what = "overall")
Data_table_10_CS <- as.matrix (Data_table_1_10, what = "classes")

write.xlsx(Data_table_10_CM, file = "Data_LDA_Model_test.xlsx", sheetName = "10PC_CM")
write.xlsx(Data_table_10_OS, file = "Data_LDA_Model_test.xlsx", sheetName = "10PC_OS", append = TRUE)
write.xlsx(Data_table_10_CS, file = "Data_LDA_Model_test.xlsx", sheetName = "10PC_CS", append = TRUE)

Data_table_20_CM <- as.matrix(Data_table_1_20)
Data_table_20_OS <- as.matrix(Data_table_1_20, what = "overall")
Data_table_20_CS <- as.matrix (Data_table_1_20, what = "classes")

write.xlsx(Data_table_20_CM, file = "Data_LDA_Model_test.xlsx", sheetName = "20PC_CM", append = TRUE)
write.xlsx(Data_table_20_OS, file = "Data_LDA_Model_test.xlsx", sheetName = "20PC_OS", append = TRUE)
write.xlsx(Data_table_20_CS, file = "Data_LDA_Model_test.xlsx", sheetName = "20PC_CS", append = TRUE)

Data_table_40_CM <- as.matrix(Data_table_1_40)
Data_table_40_OS <- as.matrix(Data_table_1_40, what = "overall")
Data_table_40_CS <- as.matrix (Data_table_1_40, what = "classes")

write.xlsx(Data_table_40_CM, file = "Data_LDA_Model_test.xlsx", sheetName = "40PC_CM", append = TRUE)
write.xlsx(Data_table_40_OS, file = "Data_LDA_Model_test.xlsx", sheetName = "40PC_OS", append = TRUE)
write.xlsx(Data_table_40_CS, file = "Data_LDA_Model_test.xlsx", sheetName = "40PC_CS", append = TRUE)

Data_table_60_CM <- as.matrix(Data_table_1_60)
Data_table_60_OS <- as.matrix(Data_table_1_60, what = "overall")
Data_table_60_CS <- as.matrix (Data_table_1_60, what = "classes")

write.xlsx(Data_table_60_CM, file = "Data_LDA_Model_test.xlsx", sheetName = "60PC_CM", append = TRUE)
write.xlsx(Data_table_60_OS, file = "Data_LDA_Model_test.xlsx", sheetName = "60PC_OS", append = TRUE)
write.xlsx(Data_table_60_CS, file = "Data_LDA_Model_test.xlsx", sheetName = "60PC_CS", append = TRUE)

Data_table_80_CM <- as.matrix(Data_table_1_80)
Data_table_80_OS <- as.matrix(Data_table_1_80, what = "overall")
Data_table_80_CS <- as.matrix (Data_table_1_80, what = "classes")

write.xlsx(Data_table_80_CM, file = "Data_LDA_Model_test.xlsx", sheetName = "80PC_CM", append = TRUE)
write.xlsx(Data_table_80_OS, file = "Data_LDA_Model_test.xlsx", sheetName = "80PC_OS", append = TRUE)
write.xlsx(Data_table_80_CS, file = "Data_LDA_Model_test.xlsx", sheetName = "80PC_CS", append = TRUE)

##Use the model statistics above to identify the model with the highest accuracy and AUC score. 
##This model will be used to complete the analysis, the model will be build at least 10 different times.
##In this example 40 PCs will be selected as the most effective model. 

#LDA Replicate 1
#Define train and test split (70% Train/30% Test) - randomly.
set.seed(11) #This is important to insure reproducibility of results, changing this number also allows for different random splits of data.
Data_40_train_size <- floor(0.7 * nrow(Data_40_DR)) #To adjust proportion of data to be assigned to training change the value (0.7) here.
set.seed(11)
Data_40_train_ind <- sample(nrow(Data_40_DR), size = Data_40_train_size)
set.seed(11)
Data_40_train_data <- as.data.frame(Data_40_DR[Data_40_train_ind,])
set.seed(11)
Data_40_test_data <- as.data.frame(Data_40_DR[-Data_40_train_ind,])

#Conduct linear discriminant analysis
set.seed (11)
Data_40_LDA <- lda(Group ~ ., data = Data_40_train_data)

#Use the LDA to make predictions
set.seed (11)
Data_40_LDA_predict <- predict(Data_40_LDA, newdata = Data_40_test_data, type = "prob")
head(Data_40_LDA_predict)

#Create and view the confusion matrix.
set.seed (11)
Data_table_1_40= table (Data_40_LDA_predict$class,Data_40_test_data$Group)
Data_table_1_40 <- confusionMatrix(Data_table_1_40)
Data_table_1_40

#Save output to file. One method is presented here but any method for saving from R can be used.
Data_table_1_CM <- as.matrix(Data_table_1_40)
Data_table_1_OS <- as.matrix(Data_table_1_40, what = "overall")
Data_table_1_CS <- as.matrix (Data_table_1_40, what = "classes")

write.xlsx(Data_table_1_CM, file = "Data_LDA_Results.xlsx", sheetName = "LDA1_CM")
write.xlsx(Data_table_1_OS, file = "Data_LDA_Results.xlsx", sheetName = "LDA1_OS", append = TRUE)
write.xlsx(Data_table_1_CS, file = "Data_LDA_Results.xlsx", sheetName = "LDA1_CS", append = TRUE)

#LDA Replicate 2
#Define train and test split (70% Train/30% Test) - randomly.
set.seed(12) #This is important to insure reproducibility of results, changing this number also allows for different random splits of data.
Data_40_train_size <- floor(0.7 * nrow(Data_40_DR)) #To adjust proportion of data to be assigned to training change the value (0.7) here.
set.seed(12)
Data_40_train_ind <- sample(nrow(Data_40_DR), size = Data_40_train_size)
set.seed(12)
Data_40_train_data <- as.data.frame(Data_40_DR[Data_40_train_ind,])
set.seed(12)
Data_40_test_data <- as.data.frame(Data_40_DR[-Data_40_train_ind,])

#Conduct linear discriminant analysis
set.seed (12)
Data_40_LDA <- lda(Group ~ ., data = Data_40_train_data)

#Use the LDA to make predictions
set.seed (12)
Data_40_LDA_predict <- predict(Data_40_LDA, newdata = Data_40_test_data, type = "prob")
head(Data_40_LDA_predict)

#Create and view the confusion matrix.
set.seed (12)
Data_table_2_40= table (Data_40_LDA_predict$class,Data_40_test_data$Group)
Data_table_2_40 <- confusionMatrix(Data_table_2_40)
Data_table_2_40

#Save output to file. One method is presented here but any method for saving from R can be used.
Data_table_2_CM <- as.matrix(Data_table_2_40)
Data_table_2_OS <- as.matrix(Data_table_2_40, what = "overall")
Data_table_2_CS <- as.matrix (Data_table_2_40, what = "classes")

write.xlsx(Data_table_2_CM, file = "Data_LDA_Results.xlsx", sheetName = "LDA2_CM", append = TRUE)
write.xlsx(Data_table_2_OS, file = "Data_LDA_Results.xlsx", sheetName = "LDA2_OS", append = TRUE)
write.xlsx(Data_table_2_CS, file = "Data_LDA_Results.xlsx", sheetName = "LDA2_CS", append = TRUE)

#LDA Replicate 3
#Define train and test split (70% Train/30% Test) - randomly.
set.seed(13) #This is important to insure reproducibility of results, changing this number also allows for different random splits of data.
Data_40_train_size <- floor(0.7 * nrow(Data_40_DR)) #To adjust proportion of data to be assigned to training change the value (0.7) here.
set.seed(13)
Data_40_train_ind <- sample(nrow(Data_40_DR), size = Data_40_train_size)
set.seed(13)
Data_40_train_data <- as.data.frame(Data_40_DR[Data_40_train_ind,])
set.seed(13)
Data_40_test_data <- as.data.frame(Data_40_DR[-Data_40_train_ind,])

#Conduct linear discriminant analysis
set.seed (13)
Data_40_LDA <- lda(Group ~ ., data = Data_40_train_data)

#Use the LDA to make predictions
set.seed (13)
Data_40_LDA_predict <- predict(Data_40_LDA, newdata = Data_40_test_data, type = "prob")
head(Data_40_LDA_predict)

#Create and view the confusion matrix.
set.seed (13)
Data_table_3_40= table (Data_40_LDA_predict$class,Data_40_test_data$Group)
Data_table_3_40 <- confusionMatrix(Data_table_3_40)
Data_table_3_40

#Save output to file. One method is presented here but any method for saving from R can be used.
Data_table_3_CM <- as.matrix(Data_table_3_40)
Data_table_3_OS <- as.matrix(Data_table_3_40, what = "overall")
Data_table_3_CS <- as.matrix (Data_table_3_40, what = "classes")

write.xlsx(Data_table_3_CM, file = "Data_LDA_Results.xlsx", sheetName = "LDA3_CM", append = TRUE)
write.xlsx(Data_table_3_OS, file = "Data_LDA_Results.xlsx", sheetName = "LDA3_OS", append = TRUE)
write.xlsx(Data_table_3_CS, file = "Data_LDA_Results.xlsx", sheetName = "LDA3_CS", append = TRUE)

#LDA Replicate 4
#Define train and test split (70% Train/30% Test) - randomly.
set.seed(14) #This is important to insure reproducibility of results, changing this number also allows for different random splits of data.
Data_40_train_size <- floor(0.7 * nrow(Data_40_DR)) #To adjust proportion of data to be assigned to training change the value (0.7) here.
set.seed(14)
Data_40_train_ind <- sample(nrow(Data_40_DR), size = Data_40_train_size)
set.seed(14)
Data_40_train_data <- as.data.frame(Data_40_DR[Data_40_train_ind,])
set.seed(14)
Data_40_test_data <- as.data.frame(Data_40_DR[-Data_40_train_ind,])

#Conduct linear discriminant analysis
set.seed (14)
Data_40_LDA <- lda(Group ~ ., data = Data_40_train_data)

#Use the LDA to make predictions
set.seed (14)
Data_40_LDA_predict <- predict(Data_40_LDA, newdata = Data_40_test_data, type = "prob")
head(Data_40_LDA_predict)

#Create and view the confusion matrix.
set.seed (14)
Data_table_4_40= table (Data_40_LDA_predict$class,Data_40_test_data$Group)
Data_table_4_40 <- confusionMatrix(Data_table_4_40)
Data_table_4_40

#Save output to file. One method is presented here but any method for saving from R can be used.
Data_table_4_CM <- as.matrix(Data_table_4_40)
Data_table_4_OS <- as.matrix(Data_table_4_40, what = "overall")
Data_table_4_CS <- as.matrix (Data_table_4_40, what = "classes")

write.xlsx(Data_table_4_CM, file = "Data_LDA_Results.xlsx", sheetName = "LDA4_CM", append = TRUE)
write.xlsx(Data_table_4_OS, file = "Data_LDA_Results.xlsx", sheetName = "LDA4_OS", append = TRUE)
write.xlsx(Data_table_4_CS, file = "Data_LDA_Results.xlsx", sheetName = "LDA4_CS", append = TRUE)

#LDA Replicate 5
#Define train and test split (70% Train/30% Test) - randomly.
set.seed(15) #This is important to insure reproducibility of results, changing this number also allows for different random splits of data.
Data_40_train_size <- floor(0.7 * nrow(Data_40_DR)) #To adjust proportion of data to be assigned to training change the value (0.7) here.
set.seed(15)
Data_40_train_ind <- sample(nrow(Data_40_DR), size = Data_40_train_size)
set.seed(15)
Data_40_train_data <- as.data.frame(Data_40_DR[Data_40_train_ind,])
set.seed(15)
Data_40_test_data <- as.data.frame(Data_40_DR[-Data_40_train_ind,])

#Conduct linear discriminant analysis
set.seed (15)
Data_40_LDA <- lda(Group ~ ., data = Data_40_train_data)

#Use the LDA to make predictions
set.seed (15)
Data_40_LDA_predict <- predict(Data_40_LDA, newdata = Data_40_test_data, type = "prob")
head(Data_40_LDA_predict)

#Create and view the confusion matrix.
set.seed (15)
Data_table_5_40= table (Data_40_LDA_predict$class,Data_40_test_data$Group)
Data_table_5_40 <- confusionMatrix(Data_table_5_40)
Data_table_5_40

#Save output to file. One method is presented here but any method for saving from R can be used.
Data_table_5_CM <- as.matrix(Data_table_5_40)
Data_table_5_OS <- as.matrix(Data_table_5_40, what = "overall")
Data_table_5_CS <- as.matrix (Data_table_5_40, what = "classes")

write.xlsx(Data_table_5_CM, file = "Data_LDA_Results.xlsx", sheetName = "LDA5_CM", append = TRUE)
write.xlsx(Data_table_5_OS, file = "Data_LDA_Results.xlsx", sheetName = "LDA5_OS", append = TRUE)
write.xlsx(Data_table_5_CS, file = "Data_LDA_Results.xlsx", sheetName = "LDA5_CS", append = TRUE)

#LDA Replicate 6
#Define train and test split (70% Train/30% Test) - randomly.
set.seed(16) #This is important to insure reproducibility of results, changing this number also allows for different random splits of data.
Data_40_train_size <- floor(0.7 * nrow(Data_40_DR)) #To adjust proportion of data to be assigned to training change the value (0.7) here.
set.seed(16)
Data_40_train_ind <- sample(nrow(Data_40_DR), size = Data_40_train_size)
set.seed(16)
Data_40_train_data <- as.data.frame(Data_40_DR[Data_40_train_ind,])
set.seed(16)
Data_40_test_data <- as.data.frame(Data_40_DR[-Data_40_train_ind,])

#Conduct linear discriminant analysis
set.seed (16)
Data_40_LDA <- lda(Group ~ ., data = Data_40_train_data)

#Use the LDA to make predictions
set.seed (16)
Data_40_LDA_predict <- predict(Data_40_LDA, newdata = Data_40_test_data, type = "prob")
head(Data_40_LDA_predict)

#Create and view the confusion matrix.
set.seed (16)
Data_table_6_40= table (Data_40_LDA_predict$class,Data_40_test_data$Group)
Data_table_6_40 <- confusionMatrix(Data_table_6_40)
Data_table_6_40

#Save output to file. One method is presented here but any method for saving from R can be used.
Data_table_6_CM <- as.matrix(Data_table_6_40)
Data_table_6_OS <- as.matrix(Data_table_6_40, what = "overall")
Data_table_6_CS <- as.matrix (Data_table_6_40, what = "classes")

write.xlsx(Data_table_6_CM, file = "Data_LDA_Results.xlsx", sheetName = "LDA6_CM", append = TRUE)
write.xlsx(Data_table_6_OS, file = "Data_LDA_Results.xlsx", sheetName = "LDA6_OS", append = TRUE)
write.xlsx(Data_table_6_CS, file = "Data_LDA_Results.xlsx", sheetName = "LDA6_CS", append = TRUE)

#LDA Replicate 7
#Define train and test split (70% Train/30% Test) - randomly.
set.seed(17) #This is important to insure reproducibility of results, changing this number also allows for different random splits of data.
Data_40_train_size <- floor(0.7 * nrow(Data_40_DR)) #To adjust proportion of data to be assigned to training change the value (0.7) here.
set.seed(17)
Data_40_train_ind <- sample(nrow(Data_40_DR), size = Data_40_train_size)
set.seed(17)
Data_40_train_data <- as.data.frame(Data_40_DR[Data_40_train_ind,])
set.seed(17)
Data_40_test_data <- as.data.frame(Data_40_DR[-Data_40_train_ind,])

#Conduct linear discriminant analysis
set.seed (17)
Data_40_LDA <- lda(Group ~ ., data = Data_40_train_data)

#Use the LDA to make predictions
set.seed (17)
Data_40_LDA_predict <- predict(Data_40_LDA, newdata = Data_40_test_data, type = "prob")
head(Data_40_LDA_predict)

#Create and view the confusion matrix.
set.seed (17)
Data_table_7_40= table (Data_40_LDA_predict$class,Data_40_test_data$Group)
Data_table_7_40 <- confusionMatrix(Data_table_7_40)
Data_table_7_40

#Save output to file. One method is presented here but any method for saving from R can be used.
Data_table_7_CM <- as.matrix(Data_table_7_40)
Data_table_7_OS <- as.matrix(Data_table_7_40, what = "overall")
Data_table_7_CS <- as.matrix (Data_table_7_40, what = "classes")

write.xlsx(Data_table_7_CM, file = "Data_LDA_Results.xlsx", sheetName = "LDA7_CM", append = TRUE)
write.xlsx(Data_table_7_OS, file = "Data_LDA_Results.xlsx", sheetName = "LDA7_OS", append = TRUE)
write.xlsx(Data_table_7_CS, file = "Data_LDA_Results.xlsx", sheetName = "LDA7_CS", append = TRUE)

#LDA Replicate 8
#Define train and test split (70% Train/30% Test) - randomly.
set.seed(18) #This is important to insure reproducibility of results, changing this number also allows for different random splits of data.
Data_40_train_size <- floor(0.7 * nrow(Data_40_DR)) #To adjust proportion of data to be assigned to training change the value (0.7) here.
set.seed(18)
Data_40_train_ind <- sample(nrow(Data_40_DR), size = Data_40_train_size)
set.seed(18)
Data_40_train_data <- as.data.frame(Data_40_DR[Data_40_train_ind,])
set.seed(18)
Data_40_test_data <- as.data.frame(Data_40_DR[-Data_40_train_ind,])

#Conduct linear discriminant analysis
set.seed (18)
Data_40_LDA <- lda(Group ~ ., data = Data_40_train_data)

#Use the LDA to make predictions
set.seed (18)
Data_40_LDA_predict <- predict(Data_40_LDA, newdata = Data_40_test_data, type = "prob")
head(Data_40_LDA_predict)

#Create and view the confusion matrix.
set.seed (18)
Data_table_8_40= table (Data_40_LDA_predict$class,Data_40_test_data$Group)
Data_table_8_40 <- confusionMatrix(Data_table_8_40)
Data_table_8_40

#Save output to file. One method is presented here but any method for saving from R can be used.
Data_table_8_CM <- as.matrix(Data_table_8_40)
Data_table_8_OS <- as.matrix(Data_table_8_40, what = "overall")
Data_table_8_CS <- as.matrix (Data_table_8_40, what = "classes")

write.xlsx(Data_table_8_CM, file = "Data_LDA_Results.xlsx", sheetName = "LDA8_CM", append = TRUE)
write.xlsx(Data_table_8_OS, file = "Data_LDA_Results.xlsx", sheetName = "LDA8_OS", append = TRUE)
write.xlsx(Data_table_8_CS, file = "Data_LDA_Results.xlsx", sheetName = "LDA8_CS", append = TRUE)

#LDA Replicate 9
#Define train and test split (70% Train/30% Test) - randomly.
set.seed(19) #This is important to insure reproducibility of results, changing this number also allows for different random splits of data.
Data_40_train_size <- floor(0.7 * nrow(Data_40_DR)) #To adjust proportion of data to be assigned to training change the value (0.7) here.
set.seed(19)
Data_40_train_ind <- sample(nrow(Data_40_DR), size = Data_40_train_size)
set.seed(19)
Data_40_train_data <- as.data.frame(Data_40_DR[Data_40_train_ind,])
set.seed(19)
Data_40_test_data <- as.data.frame(Data_40_DR[-Data_40_train_ind,])

#Conduct linear discriminant analysis
set.seed (19)
Data_40_LDA <- lda(Group ~ ., data = Data_40_train_data)

#Use the LDA to make predictions
set.seed (19)
Data_40_LDA_predict <- predict(Data_40_LDA, newdata = Data_40_test_data, type = "prob")
head(Data_40_LDA_predict)

#Create and view the confusion matrix.
set.seed (19)
Data_table_9_40= table (Data_40_LDA_predict$class,Data_40_test_data$Group)
Data_table_9_40 <- confusionMatrix(Data_table_9_40)
Data_table_9_40

#Save output to file. One method is presented here but any method for saving from R can be used.
Data_table_9_CM <- as.matrix(Data_table_9_40)
Data_table_9_OS <- as.matrix(Data_table_9_40, what = "overall")
Data_table_9_CS <- as.matrix (Data_table_9_40, what = "classes")

write.xlsx(Data_table_9_CM, file = "Data_LDA_Results.xlsx", sheetName = "LDA9_CM", append = TRUE)
write.xlsx(Data_table_9_OS, file = "Data_LDA_Results.xlsx", sheetName = "LDA9_OS", append = TRUE)
write.xlsx(Data_table_9_CS, file = "Data_LDA_Results.xlsx", sheetName = "LDA9_CS", append = TRUE)

#LDA Replicate 10
#Define train and test split (70% Train/30% Test) - randomly.
set.seed(20) #This is important to insure reproducibility of results, changing this number also allows for different random splits of data.
Data_40_train_size <- floor(0.7 * nrow(Data_40_DR)) #To adjust proportion of data to be assigned to training change the value (0.7) here.
set.seed(20)
Data_40_train_ind <- sample(nrow(Data_40_DR), size = Data_40_train_size)
set.seed(20)
Data_40_train_data <- as.data.frame(Data_40_DR[Data_40_train_ind,])
set.seed(20)
Data_40_test_data <- as.data.frame(Data_40_DR[-Data_40_train_ind,])

#Conduct linear discriminant analysis
set.seed (20)
Data_40_LDA <- lda(Group ~ ., data = Data_40_train_data)

#Use the LDA to make predictions
set.seed (20)
Data_40_LDA_predict <- predict(Data_40_LDA, newdata = Data_40_test_data, type = "prob")
head(Data_40_LDA_predict)

#Create and view the confusion matrix.
set.seed (20)
Data_table_10_40= table (Data_40_LDA_predict$class,Data_40_test_data$Group)
Data_table_10_40 <- confusionMatrix(Data_table_10_40)
Data_table_10_40

#Save output to file. One method is presented here but any method for saving from R can be used.
Data_table_10_CM <- as.matrix(Data_table_10_40)
Data_table_10_OS <- as.matrix(Data_table_10_40, what = "overall")
Data_table_10_CS <- as.matrix (Data_table_10_40, what = "classes")

write.xlsx(Data_table_10_CM, file = "Data_LDA_Results.xlsx", sheetName = "LDA10_CM", append = TRUE)
write.xlsx(Data_table_10_OS, file = "Data_LDA_Results.xlsx", sheetName = "LDA10_OS", append = TRUE)
write.xlsx(Data_table_10_CS, file = "Data_LDA_Results.xlsx", sheetName = "LDA10_CS", append = TRUE)

##Analysis using LDA as a classifier is now complete. The output file contains model statistics and confusion matrix for further use.

##Random forest (RF) is going to be used as a classifier.
##For classification models the data will be split randomly into 70% training and 30% test data. These proportions can be adjusted to suit data.
##The RF should be repeated at least 10x with different train and test data.

##Use the first set of train/test data determine optimal number of PCs to use in RF.This should be completed for the data sets with differing numbers of principle comonents compared above.
##For RF the tree number (ntree) and number of variables randomly sampled as candidates at each split (mtry) must also be optimised. 
##PC number, ntree and mtry will be optimised by cross vallidation using the caret package.
##This example will use manual tuning in caret to tune parameters, the mtry and ntree number will be depicted by the user. 
##Other methods including grid and random searching can also be used in the caret package.


##Create custom carot random forest for tuning mtry and ntree
customRF <- list(type = "Classification", #Model type that will be used is classification.
                 library = "randomForest", #Other models can be used in caret so here specify which model is to be used.
                 loop = NULL)

customRF$parameters <- data.frame(parameter = c("mtry", "ntree"),
                                  class = rep("numeric", 2),
                                  label = c("mtry", "ntree"))

customRF$grid <- function(x, y, len = NULL, search = "grid") {}

customRF$fit <- function(x, y, wts, param, lev, last, weights, classProbs) {
  randomForest(x, y,
               mtry = param$mtry, #mtry will be depicted by the object mtry which will be created below, with user depicted mtry values.
               ntree=param$ntree) #ntree will be depicted by the object mtry which will be created below, with user depcited values.
}

#Predict label
customRF$predict <- function(modelFit, newdata, preProc = NULL, submodels = NULL)
  predict(modelFit, newdata)

#Predict prob
customRF$prob <- function(modelFit, newdata, preProc = NULL, submodels = NULL)
  predict(modelFit, newdata, type = "prob")

customRF$sort <- function(x) x[order(x[,1]),]
customRF$levels <- function(x) x$classes

##Create control function for training with 10 folds and keep 3 folds for training.
control <- trainControl(method="repeatedcv", #This uses repeated cross vallidation for model tuning.
                        number=10, #Here 10 folds are being used, this number can be increased to suit data.
                        repeats=3,#Here 3 replicates are being used, this number can be increased to suit data.
                        allowParallel = TRUE, sampling = "down") #For unbalanced data, over or undersampling may be used to reduce negative impacts of the unbalance. For over sampling 'sampling = "up"' should be used, if no under/over sampling is required this can be removed.


##The control and custom random forest are now ready to use with the dataset.

##Use the first set of train/test data determine optimal number of PCs to use in RF.This should be completed for the data sets with differing numbers of principle comonents compared above.

#Testing 10 PCs

#create tunegrid with various mtry and ntree values for tuning the model.
#mtry should be a value ranging from 2 - total number of variables in the data set (will be different for each of the reduced datasets).
#ntree numbers can be experimented with to find optimal. 
tunegrid_10 <- expand.grid(.mtry = (c(2,4,6,8,10)), .ntree = c(500,1000,1500,2000,4000,6000,8000))

#Run caret with 10 PCs to find optimal parameters. 
set.seed (1)
Data_train_size <- floor(0.7 * nrow(Data_10_DR))
set.seed (1)
Data_train_ind <- sample(nrow(Data_10_DR), size = Data_train_size)
set.seed (1)
Data_train_data <- as.data.frame(Data_10_DR[Data_train_ind,])
set.seed (1)
Data_test_data <- as.data.frame(Data_10_DR[-Data_train_ind,])

Data_Caret_10 <- train(Group~., data=Data_train_data, 
                      method=customRF, 
                      metric='Accuracy', 
                      tuneGrid=tunegrid_10, 
                      trControl=control)

plot (Data_Caret_10)
print (Data_Caret_10)

#Testing 20 PCs

#create tunegrid with various mtry and ntree values for tuning the model.
#mtry should be a value ranging from 2 - total number of variables in the data set (will be different for each of the reduced datasets).
#ntree numbers can be experimented with to find optimal. 
tunegrid_20 <- expand.grid(.mtry = (c(2,5,10,15,20)), .ntree = c(500,2000,1500,2000,4000,6000,8000))

#Run caret with 20 PCs to find optimal parameters. 
set.seed (1)
Data_train_size <- floor(0.7 * nrow(Data_20_DR))
set.seed (1)
Data_train_ind <- sample(nrow(Data_20_DR), size = Data_train_size)
set.seed (1)
Data_train_data <- as.data.frame(Data_20_DR[Data_train_ind,])
set.seed (1)
Data_test_data <- as.data.frame(Data_20_DR[-Data_train_ind,])

Data_Caret_20 <- train(Group~., data=Data_train_data, 
                       method=customRF, 
                       metric='Accuracy', 
                       tuneGrid=tunegrid_20, 
                       trControl=control)

plot (Data_Caret_20)
print (Data_Caret_20)

#Testing 40 PCs

#create tunegrid with various mtry and ntree values for tuning the model.
#mtry should be a value ranging from 2 - total number of variables in the data set (will be different for each of the reduced datasets).
#ntree numbers can be experimented with to find optimal. 
tunegrid_40 <- expand.grid(.mtry = (c(10,20,30,40)), .ntree = c(500,4000,1500,4000,4000,6000,8000))

#Run caret with 40 PCs to find optimal parameters. 
set.seed (1)
Data_train_size <- floor(0.7 * nrow(Data_40_DR))
set.seed (1)
Data_train_ind <- sample(nrow(Data_40_DR), size = Data_train_size)
set.seed (1)
Data_train_data <- as.data.frame(Data_40_DR[Data_train_ind,])
set.seed (1)
Data_test_data <- as.data.frame(Data_40_DR[-Data_train_ind,])

Data_Caret_40 <- train(Group~., data=Data_train_data, 
                       method=customRF, 
                       metric='Accuracy', 
                       tuneGrid=tunegrid_40, 
                       trControl=control)

plot (Data_Caret_40)
print (Data_Caret_40)

#Testing 60 PCs

#create tunegrid with various mtry and ntree values for tuning the model.
#mtry should be a value ranging from 2 - total number of variables in the data set (will be different for each of the reduced datasets).
#ntree numbers can be experimented with to find optimal. 
tunegrid_60 <- expand.grid(.mtry = (c(10,20,30,40,50,60)), .ntree = c(500,6000,1500,6000,6000,6000,8000))

#Run caret with 60 PCs to find optimal parameters. 
set.seed (1)
Data_train_size <- floor(0.7 * nrow(Data_60_DR))
set.seed (1)
Data_train_ind <- sample(nrow(Data_60_DR), size = Data_train_size)
set.seed (1)
Data_train_data <- as.data.frame(Data_60_DR[Data_train_ind,])
set.seed (1)
Data_test_data <- as.data.frame(Data_60_DR[-Data_train_ind,])

Data_Caret_60 <- train(Group~., data=Data_train_data, 
                       method=customRF, 
                       metric='Accuracy', 
                       tuneGrid=tunegrid_60, 
                       trControl=control)

plot (Data_Caret_60)
print (Data_Caret_60)

#Testing 80 PCs

#create tunegrid with various mtry and ntree values for tuning the model.
#mtry should be a value ranging from 2 - total number of variables in the data set (will be different for each of the reduced datasets).
#ntree numbers can be experimented with to find optimal. 
tunegrid_80 <- expand.grid(.mtry = (c(10,20,30,40,50,60,70,80)), .ntree = c(500,8000,1500,8000,8000,8000,8000))

#Run caret with 80 PCs to find optimal parameters. 
set.seed (1)
Data_train_size <- floor(0.7 * nrow(Data_80_DR))
set.seed (1)
Data_train_ind <- sample(nrow(Data_80_DR), size = Data_train_size)
set.seed (1)
Data_train_data <- as.data.frame(Data_80_DR[Data_train_ind,])
set.seed (1)
Data_test_data <- as.data.frame(Data_80_DR[-Data_train_ind,])

Data_Caret_80 <- train(Group~., data=Data_train_data, 
                       method=customRF, 
                       metric='Accuracy', 
                       tuneGrid=tunegrid_80,
                       trControl=control)

plot (Data_Caret_80)
print (Data_Caret_80)

##Caret plots and outputs can be saved for future use. 

##Use the caret output above to identify the model with the highest accuracy. Use the mtry and ntree values identified by caret.
##This model will be used to complete the analysis, the model will be build at least 10 different times.
##In this example 20 PCs will be selected as the most effective model. 


#Random Forest Rep 1
#Define train and test split (70% Train/30% Test) - randomly.
set.seed (11) #This is important to insure reproducibility of results, changing this number also allows for different random splits of data.
Data_train_size <- floor(0.8 * nrow(Data_20_DR)) #To adjust proportion of data to be assigned to training change the value (0.7) here.
set.seed (11)
Data_train_ind <- sample(nrow(Data_20_DR), size = Data_train_size)
set.seed (11)
Data_train_data <- as.data.frame(Data_20_DR[Data_train_ind,])
set.seed (11)
Data_test_data <- as.data.frame(Data_20_DR[-Data_train_ind,])

#Balance the data using over or under sampling. If classes are balanced this step can be skipped.
sum (Data_train_data$Group == 1)
sum (Data_train_data$Group == 2)

Data_train_data <- downSample(Data_train_data, Data_train_data$Group, list = FALSE, yname = "Group") #To use random over sampling change command to upSample.
Data_train_data <- Data_train_data [,-22]

sum (Data_train_data$Group == 1)
sum (Data_train_data$Group == 2)

#Conduct random forest analysis.
set.seed (11)
Data_RF1 <- randomForest(Group ~ ., data=Data_train_data, mtry =2 , ntree = 2000)
print (Data_RF1)

#Use the random forest to make predictions.
set.seed (11)
pred_Data_RF1 = predict (Data_RF1, newdata = Data_test_data [,-21])

#Create and view the confusion matrix.
pred_Data_RF1 = table (Data_test_data[,21], pred_Data_RF1)
pred_Data_RF1 <- confusionMatrix(pred_Data_RF1)

#Save output to file. One method is presented here but any method for saving from R can be used.
Data_table_1_CM <- as.matrix(pred_Data_RF1)
Data_table_1_OS <- as.matrix(pred_Data_RF1, what = "overall")
Data_table_1_CS <- as.matrix (pred_Data_RF1, what = "classes")

write.xlsx(Data_table_1_CM, file = "Data_RF_Results.xlsx", sheetName = "RF1_CM")
write.xlsx(Data_table_1_OS, file = "Data_RF_Results.xlsx", sheetName = "RF1_OS", append = TRUE)
write.xlsx(Data_table_1_CS, file = "Data_RF_Results.xlsx", sheetName = "RF1_CS", append = TRUE)

#Random Forest Rep 2
#Define train and test split (70% Train/30% Test) - randomly.
set.seed (12) #This is important to insure reproducibility of results, changing this number also allows for different random splits of data.
Data_train_size <- floor(0.8 * nrow(Data_20_DR)) #To adjust proportion of data to be assigned to training change the value (0.7) here.
set.seed (12)
Data_train_ind <- sample(nrow(Data_20_DR), size = Data_train_size)
set.seed (12)
Data_train_data <- as.data.frame(Data_20_DR[Data_train_ind,])
set.seed (12)
Data_test_data <- as.data.frame(Data_20_DR[-Data_train_ind,])

#Balance the data using over or under sampling. If classes are balanced this step can be skipped.
sum (Data_train_data$Group == 1)
sum (Data_train_data$Group == 2)

Data_train_data <- downSample(Data_train_data, Data_train_data$Group, list = FALSE, yname = "Group") #To use random over sampling change command to upSample.
Data_train_data <- Data_train_data [,-22]

sum (Data_train_data$Group == 1)
sum (Data_train_data$Group == 2)

#Conduct random forest analysis.
set.seed (12)
Data_RF2 <- randomForest(Group ~ ., data=Data_train_data, mtry =2 , ntree = 2000)
print (Data_RF2)

#Use the random forest to make predictions.
set.seed (12)
pred_Data_RF2 = predict (Data_RF2, newdata = Data_test_data [,-21])

#Create and view the confusion matrix.
pred_Data_RF2 = table (Data_test_data[,21], pred_Data_RF2)
pred_Data_RF2 <- confusionMatrix(pred_Data_RF2)

#Save output to file. One method is presented here but any method for saving from R can be used.
Data_table_2_CM <- as.matrix(pred_Data_RF2)
Data_table_2_OS <- as.matrix(pred_Data_RF2, what = "overall")
Data_table_2_CS <- as.matrix (pred_Data_RF2, what = "classes")

write.xlsx(Data_table_2_CM, file = "Data_RF_Results.xlsx", sheetName = "RF2_CM", append = TRUE)
write.xlsx(Data_table_2_OS, file = "Data_RF_Results.xlsx", sheetName = "RF2_OS", append = TRUE)
write.xlsx(Data_table_2_CS, file = "Data_RF_Results.xlsx", sheetName = "RF2_CS", append = TRUE)

#Random Forest Rep 3
#Define train and test split (70% Train/30% Test) - randomly.
set.seed (13) #This is important to insure reproducibility of results, changing this number also allows for different random splits of data.
Data_train_size <- floor(0.8 * nrow(Data_20_DR)) #To adjust proportion of data to be assigned to training change the value (0.7) here.
set.seed (13)
Data_train_ind <- sample(nrow(Data_20_DR), size = Data_train_size)
set.seed (13)
Data_train_data <- as.data.frame(Data_20_DR[Data_train_ind,])
set.seed (13)
Data_test_data <- as.data.frame(Data_20_DR[-Data_train_ind,])

#Balance the data using over or under sampling. If classes are balanced this step can be skipped.
sum (Data_train_data$Group == 1)
sum (Data_train_data$Group == 2)

Data_train_data <- downSample(Data_train_data, Data_train_data$Group, list = FALSE, yname = "Group") #To use random over sampling change command to upSample.
Data_train_data <- Data_train_data [,-22]

sum (Data_train_data$Group == 1)
sum (Data_train_data$Group == 2)

#Conduct random forest analysis.
set.seed (13)
Data_RF3 <- randomForest(Group ~ ., data=Data_train_data, mtry =2 , ntree = 2000)
print (Data_RF3)

#Use the random forest to make predictions.
set.seed (13)
pred_Data_RF3 = predict (Data_RF3, newdata = Data_test_data [,-21])

#Create and view the confusion matrix.
pred_Data_RF3 = table (Data_test_data[,21], pred_Data_RF3)
pred_Data_RF3 <- confusionMatrix(pred_Data_RF3)

#Save output to file. One method is presented here but any method for saving from R can be used.
Data_table_3_CM <- as.matrix(pred_Data_RF3)
Data_table_3_OS <- as.matrix(pred_Data_RF3, what = "overall")
Data_table_3_CS <- as.matrix (pred_Data_RF3, what = "classes")

write.xlsx(Data_table_3_CM, file = "Data_RF_Results.xlsx", sheetName = "RF3_CM", append = TRUE)
write.xlsx(Data_table_3_OS, file = "Data_RF_Results.xlsx", sheetName = "RF3_OS", append = TRUE)
write.xlsx(Data_table_3_CS, file = "Data_RF_Results.xlsx", sheetName = "RF3_CS", append = TRUE)

#Random Forest Rep 4
#Define train and test split (70% Train/40% Test) - randomly.
set.seed (14) #This is important to insure reproducibility of results, changing this number also allows for different random splits of data.
Data_train_size <- floor(0.8 * nrow(Data_20_DR)) #To adjust proportion of data to be assigned to training change the value (0.7) here.
set.seed (14)
Data_train_ind <- sample(nrow(Data_20_DR), size = Data_train_size)
set.seed (14)
Data_train_data <- as.data.frame(Data_20_DR[Data_train_ind,])
set.seed (14)
Data_test_data <- as.data.frame(Data_20_DR[-Data_train_ind,])

#Balance the data using over or under sampling. If classes are balanced this step can be skipped.
sum (Data_train_data$Group == 1)
sum (Data_train_data$Group == 2)

Data_train_data <- downSample(Data_train_data, Data_train_data$Group, list = FALSE, yname = "Group") #To use random over sampling change command to upSample.
Data_train_data <- Data_train_data [,-22]

sum (Data_train_data$Group == 1)
sum (Data_train_data$Group == 2)

#Conduct random forest analysis.
set.seed (14)
Data_RF4 <- randomForest(Group ~ ., data=Data_train_data, mtry =2 , ntree = 2000)
print (Data_RF4)

#Use the random forest to make predictions.
set.seed (14)
pred_Data_RF4 = predict (Data_RF4, newdata = Data_test_data [,-21])

#Create and view the confusion matrix.
pred_Data_RF4 = table (Data_test_data[,21], pred_Data_RF4)
pred_Data_RF4 <- confusionMatrix(pred_Data_RF4)

#Save output to file. One method is presented here but any method for saving from R can be used.
Data_table_4_CM <- as.matrix(pred_Data_RF4)
Data_table_4_OS <- as.matrix(pred_Data_RF4, what = "overall")
Data_table_4_CS <- as.matrix (pred_Data_RF4, what = "classes")

write.xlsx(Data_table_4_CM, file = "Data_RF_Results.xlsx", sheetName = "RF4_CM", append = TRUE)
write.xlsx(Data_table_4_OS, file = "Data_RF_Results.xlsx", sheetName = "RF4_OS", append = TRUE)
write.xlsx(Data_table_4_CS, file = "Data_RF_Results.xlsx", sheetName = "RF4_CS", append = TRUE)

#Random Forest Rep 5
#Define train and test split (70% Train/50% Test) - randomly.
set.seed (15) #This is important to insure reproducibility of results, changing this number also allows for different random splits of data.
Data_train_size <- floor(0.8 * nrow(Data_20_DR)) #To adjust proportion of data to be assigned to training change the value (0.7) here.
set.seed (15)
Data_train_ind <- sample(nrow(Data_20_DR), size = Data_train_size)
set.seed (15)
Data_train_data <- as.data.frame(Data_20_DR[Data_train_ind,])
set.seed (15)
Data_test_data <- as.data.frame(Data_20_DR[-Data_train_ind,])

#Balance the data using over or under sampling. If classes are balanced this step can be skipped.
sum (Data_train_data$Group == 1)
sum (Data_train_data$Group == 2)

Data_train_data <- downSample(Data_train_data, Data_train_data$Group, list = FALSE, yname = "Group") #To use random over sampling change command to upSample.
Data_train_data <- Data_train_data [,-22]

sum (Data_train_data$Group == 1)
sum (Data_train_data$Group == 2)

#Conduct random forest analysis.
set.seed (15)
Data_RF5 <- randomForest(Group ~ ., data=Data_train_data, mtry =2 , ntree = 2000)
print (Data_RF5)

#Use the random forest to make predictions.
set.seed (15)
pred_Data_RF5 = predict (Data_RF5, newdata = Data_test_data [,-21])

#Create and view the confusion matrix.
pred_Data_RF5 = table (Data_test_data[,21], pred_Data_RF5)
pred_Data_RF5 <- confusionMatrix(pred_Data_RF5)

#Save output to file. One method is presented here but any method for saving from R can be used.
Data_table_5_CM <- as.matrix(pred_Data_RF5)
Data_table_5_OS <- as.matrix(pred_Data_RF5, what = "overall")
Data_table_5_CS <- as.matrix (pred_Data_RF5, what = "classes")

write.xlsx(Data_table_5_CM, file = "Data_RF_Results.xlsx", sheetName = "RF5_CM", append = TRUE)
write.xlsx(Data_table_5_OS, file = "Data_RF_Results.xlsx", sheetName = "RF5_OS", append = TRUE)
write.xlsx(Data_table_5_CS, file = "Data_RF_Results.xlsx", sheetName = "RF5_CS", append = TRUE)

#Random Forest Rep 6
#Define train and test split (70% Train/60% Test) - randomly.
set.seed (16) #This is important to insure reproducibility of results, changing this number also allows for different random splits of data.
Data_train_size <- floor(0.8 * nrow(Data_20_DR)) #To adjust proportion of data to be assigned to training change the value (0.7) here.
set.seed (16)
Data_train_ind <- sample(nrow(Data_20_DR), size = Data_train_size)
set.seed (16)
Data_train_data <- as.data.frame(Data_20_DR[Data_train_ind,])
set.seed (16)
Data_test_data <- as.data.frame(Data_20_DR[-Data_train_ind,])

#Balance the data using over or under sampling. If classes are balanced this step can be skipped.
sum (Data_train_data$Group == 1)
sum (Data_train_data$Group == 2)

Data_train_data <- downSample(Data_train_data, Data_train_data$Group, list = FALSE, yname = "Group") #To use random over sampling change command to upSample.
Data_train_data <- Data_train_data [,-22]

sum (Data_train_data$Group == 1)
sum (Data_train_data$Group == 2)

#Conduct random forest analysis.
set.seed (16)
Data_RF6 <- randomForest(Group ~ ., data=Data_train_data, mtry =2 , ntree = 2000)
print (Data_RF6)

#Use the random forest to make predictions.
set.seed (16)
pred_Data_RF6 = predict (Data_RF6, newdata = Data_test_data [,-21])

#Create and view the confusion matrix.
pred_Data_RF6 = table (Data_test_data[,21], pred_Data_RF6)
pred_Data_RF6 <- confusionMatrix(pred_Data_RF6)

#Save output to file. One method is presented here but any method for saving from R can be used.
Data_table_6_CM <- as.matrix(pred_Data_RF6)
Data_table_6_OS <- as.matrix(pred_Data_RF6, what = "overall")
Data_table_6_CS <- as.matrix (pred_Data_RF6, what = "classes")

write.xlsx(Data_table_6_CM, file = "Data_RF_Results.xlsx", sheetName = "RF6_CM")
write.xlsx(Data_table_6_OS, file = "Data_RF_Results.xlsx", sheetName = "RF6_OS", append = TRUE)
write.xlsx(Data_table_6_CS, file = "Data_RF_Results.xlsx", sheetName = "RF6_CS", append = TRUE)

#Random Forest Rep 7
#Define train and test split (70% Train/70% Test) - randomly.
set.seed (17) #This is important to insure reproducibility of results, changing this number also allows for different random splits of data.
Data_train_size <- floor(0.8 * nrow(Data_20_DR)) #To adjust proportion of data to be assigned to training change the value (0.7) here.
set.seed (17)
Data_train_ind <- sample(nrow(Data_20_DR), size = Data_train_size)
set.seed (17)
Data_train_data <- as.data.frame(Data_20_DR[Data_train_ind,])
set.seed (17)
Data_test_data <- as.data.frame(Data_20_DR[-Data_train_ind,])

#Balance the data using over or under sampling. If classes are balanced this step can be skipped.
sum (Data_train_data$Group == 1)
sum (Data_train_data$Group == 2)

Data_train_data <- downSample(Data_train_data, Data_train_data$Group, list = FALSE, yname = "Group") #To use random over sampling change command to upSample.
Data_train_data <- Data_train_data [,-22]

sum (Data_train_data$Group == 1)
sum (Data_train_data$Group == 2)

#Conduct random forest analysis.
set.seed (17)
Data_RF7 <- randomForest(Group ~ ., data=Data_train_data, mtry =2 , ntree = 2000)
print (Data_RF7)

#Use the random forest to make predictions.
set.seed (17)
pred_Data_RF7 = predict (Data_RF7, newdata = Data_test_data [,-21])

#Create and view the confusion matrix.
pred_Data_RF7 = table (Data_test_data[,21], pred_Data_RF7)
pred_Data_RF7 <- confusionMatrix(pred_Data_RF7)

#Save output to file. One method is presented here but any method for saving from R can be used.
Data_table_7_CM <- as.matrix(pred_Data_RF7)
Data_table_7_OS <- as.matrix(pred_Data_RF7, what = "overall")
Data_table_7_CS <- as.matrix (pred_Data_RF7, what = "classes")

write.xlsx(Data_table_7_CM, file = "Data_RF_Results.xlsx", sheetName = "RF7_CM", append = TRUE)
write.xlsx(Data_table_7_OS, file = "Data_RF_Results.xlsx", sheetName = "RF7_OS", append = TRUE)
write.xlsx(Data_table_7_CS, file = "Data_RF_Results.xlsx", sheetName = "RF7_CS", append = TRUE)

#Random Forest Rep 8
#Define train and test split (70% Train/80% Test) - randomly.
set.seed (18) #This is important to insure reproducibility of results, changing this number also allows for different random splits of data.
Data_train_size <- floor(0.8 * nrow(Data_20_DR)) #To adjust proportion of data to be assigned to training change the value (0.7) here.
set.seed (18)
Data_train_ind <- sample(nrow(Data_20_DR), size = Data_train_size)
set.seed (18)
Data_train_data <- as.data.frame(Data_20_DR[Data_train_ind,])
set.seed (18)
Data_test_data <- as.data.frame(Data_20_DR[-Data_train_ind,])

#Balance the data using over or under sampling. If classes are balanced this step can be skipped.
sum (Data_train_data$Group == 1)
sum (Data_train_data$Group == 2)

Data_train_data <- downSample(Data_train_data, Data_train_data$Group, list = FALSE, yname = "Group") #To use random over sampling change command to upSample.
Data_train_data <- Data_train_data [,-22]

sum (Data_train_data$Group == 1)
sum (Data_train_data$Group == 2)

#Conduct random forest analysis.
set.seed (18)
Data_RF8 <- randomForest(Group ~ ., data=Data_train_data, mtry =2 , ntree = 2000)
print (Data_RF8)

#Use the random forest to make predictions.
set.seed (18)
pred_Data_RF8 = predict (Data_RF8, newdata = Data_test_data [,-21])

#Create and view the confusion matrix.
pred_Data_RF8 = table (Data_test_data[,21], pred_Data_RF8)
pred_Data_RF8 <- confusionMatrix(pred_Data_RF8)

#Save output to file. One method is presented here but any method for saving from R can be used.
Data_table_8_CM <- as.matrix(pred_Data_RF8)
Data_table_8_OS <- as.matrix(pred_Data_RF8, what = "overall")
Data_table_8_CS <- as.matrix (pred_Data_RF8, what = "classes")

write.xlsx(Data_table_8_CM, file = "Data_RF_Results.xlsx", sheetName = "RF8_CM", append = TRUE)
write.xlsx(Data_table_8_OS, file = "Data_RF_Results.xlsx", sheetName = "RF8_OS", append = TRUE)
write.xlsx(Data_table_8_CS, file = "Data_RF_Results.xlsx", sheetName = "RF8_CS", append = TRUE)

#Random Forest Rep 9
#Define train and test split (70% Train/90% Test) - randomly.
set.seed (19) #This is important to insure reproducibility of results, changing this number also allows for different random splits of data.
Data_train_size <- floor(0.8 * nrow(Data_20_DR)) #To adjust proportion of data to be assigned to training change the value (0.7) here.
set.seed (19)
Data_train_ind <- sample(nrow(Data_20_DR), size = Data_train_size)
set.seed (19)
Data_train_data <- as.data.frame(Data_20_DR[Data_train_ind,])
set.seed (19)
Data_test_data <- as.data.frame(Data_20_DR[-Data_train_ind,])

#Balance the data using over or under sampling. If classes are balanced this step can be skipped.
sum (Data_train_data$Group == 1)
sum (Data_train_data$Group == 2)

Data_train_data <- downSample(Data_train_data, Data_train_data$Group, list = FALSE, yname = "Group") #To use random over sampling change command to upSample.
Data_train_data <- Data_train_data [,-22]

sum (Data_train_data$Group == 1)
sum (Data_train_data$Group == 2)

#Conduct random forest analysis.
set.seed (19)
Data_RF9 <- randomForest(Group ~ ., data=Data_train_data, mtry =2 , ntree = 2000)
print (Data_RF9)

#Use the random forest to make predictions.
set.seed (19)
pred_Data_RF9 = predict (Data_RF9, newdata = Data_test_data [,-21])

#Create and view the confusion matrix.
pred_Data_RF9 = table (Data_test_data[,21], pred_Data_RF9)
pred_Data_RF9 <- confusionMatrix(pred_Data_RF9)

#Save output to file. One method is presented here but any method for saving from R can be used.
Data_table_9_CM <- as.matrix(pred_Data_RF9)
Data_table_9_OS <- as.matrix(pred_Data_RF9, what = "overall")
Data_table_9_CS <- as.matrix (pred_Data_RF9, what = "classes")

write.xlsx(Data_table_9_CM, file = "Data_RF_Results.xlsx", sheetName = "RF9_CM", append = TRUE)
write.xlsx(Data_table_9_OS, file = "Data_RF_Results.xlsx", sheetName = "RF9_OS", append = TRUE)
write.xlsx(Data_table_9_CS, file = "Data_RF_Results.xlsx", sheetName = "RF9_CS", append = TRUE)

#Random Forest Rep 10
#Define train and test split (70% Train/100% Test) - randomly.
set.seed (20) #This is important to insure reproducibility of results, changing this number also allows for different random splits of data.
Data_train_size <- floor(0.8 * nrow(Data_20_DR)) #To adjust proportion of data to be assigned to training change the value (0.7) here.
set.seed (20)
Data_train_ind <- sample(nrow(Data_20_DR), size = Data_train_size)
set.seed (20)
Data_train_data <- as.data.frame(Data_20_DR[Data_train_ind,])
set.seed (20)
Data_test_data <- as.data.frame(Data_20_DR[-Data_train_ind,])

#Balance the data using over or under sampling. If classes are balanced this step can be skipped.
sum (Data_train_data$Group == 1)
sum (Data_train_data$Group == 2)

Data_train_data <- downSample(Data_train_data, Data_train_data$Group, list = FALSE, yname = "Group") #To use random over sampling change command to upSample.
Data_train_data <- Data_train_data [,-22]

sum (Data_train_data$Group == 1)
sum (Data_train_data$Group == 2)

#Conduct random forest analysis.
set.seed (20)
Data_RF10 <- randomForest(Group ~ ., data=Data_train_data, mtry =2 , ntree = 2000)
print (Data_RF10)

#Use the random forest to make predictions.
set.seed (20)
pred_Data_RF10 = predict (Data_RF10, newdata = Data_test_data [,-21])

#Create and view the confusion matrix.
pred_Data_RF10 = table (Data_test_data[,21], pred_Data_RF10)
pred_Data_RF10 <- confusionMatrix(pred_Data_RF10)

#Save output to file. One method is presented here but any method for saving from R can be used.
Data_table_10_CM <- as.matrix(pred_Data_RF10)
Data_table_10_OS <- as.matrix(pred_Data_RF10, what = "overall")
Data_table_10_CS <- as.matrix (pred_Data_RF10, what = "classes")

write.xlsx(Data_table_10_CM, file = "Data_RF_Results.xlsx", sheetName = "RF10_CM", append = TRUE)
write.xlsx(Data_table_10_OS, file = "Data_RF_Results.xlsx", sheetName = "RF10_OS", append = TRUE)
write.xlsx(Data_table_10_CS, file = "Data_RF_Results.xlsx", sheetName = "RF10_CS", append = TRUE)

##Analysis using random forest as a classifier is now complete. The output file contains model statistics and confusion matrix for further use.
